# Intracranial recordings demonstrate medial temporal lobe engagement in visual search in humans

**DOI:** 10.1101/2020.02.29.971341

**Authors:** S. J. Katarina Slama, Richard Jimenez, Sujayam Saha, David King-Stephens, Kenneth D. Laxer, Peter B. Weber, Tor Endestad, Pål G. Larsson, Anne-Kristin Solbakk, Jack J. Lin, Robert T. Knight

## Abstract

Visual search is a fundamental human behavior, which has been proposed to include two component processes: inefficient search (Search) and efficient search (Pop-out). According to extant research, these two processes map onto two separable neural systems located in the frontal and parietal association cortices. In the present study, we use intracranial recordings from 23 participants to delineate the neural correlates of Search and Pop-out with an unprecedented combination of spatiotemporal resolution and coverage across cortical and subcortical structures. First, we demonstrate a role for the medial temporal lobe in visual search, on par with engagement in frontal and parietal association cortex. Second, we show a gradient of increasing engagement over anatomical space from dorsal to ventral lateral frontal cortex. Third, we confirm previous work demonstrating nearly complete overlap in neural engagement across cortical regions in Search and Pop-out. We further demonstrate Pop-out selectivity manifesting as activity increase in Pop-out as compared to Search in a distributed set of sites including frontal cortex. This result is at odds with the view that Pop-out is implemented in low-level visual cortex or parietal cortex alone. Finally, we affirm a central role for the right lateral frontal cortex in Search.

## Introduction

Visual search is ubiquitous in everyday life, and is deployed in everything from driving to reading to airport security screening. Impairments in visual search ability have been documented in numerous brain diseases including Alzheimer’s disease, Parkinson’s disease, stroke, schizophrenia, and ADHD. Visual search is widely considered a classical attention behavior (Treisman & Gelade,1980; Wolfe, 2014; 2018). Accordingly, some search processes are thought to require deliberate allocation of attention to individual putative search targets in a serial fashion (inefficient search, hereafter referred to as Search), with response times (RTs) increasing with the number of distractors. In other search processes, attention is thought to be automatically captured by a salient item (efficient search, hereafter referred to as Pop-out), rendering search times independent of the number of distractors. Pop-out is thought to rely on preattentive visual perceptual mechanisms which scan the entire visual field in parallel (Treisman & Gelade, 1980; Julesz, 1981; Wolfe, 2018; but see Nakayama & Martini, 2011). Visual search experiments are viewed as well-parameterized *attention* tasks. Depending on the design of the specific experiment, Search and Pop-out are often viewed a eliciting attentive and pre-attentive processes, respectively.

### Putative medial temporal lobe engagement in Search and Pop-out

The dominant model for the neural substrates of visual attention derives from human fMRI and PET studies as well as single-neuron research in non-human primates (Corbetta & Shulman, 2002). According to this framework, a bilateral dorsal network (the dorsal attention network, DAN) supports top-down attention, as deployed in Search. A right-lateralized ventral attention network (the ventral attention network, VAN) is implicated in bottom-up attention, as deployed in Pop-out (Corbetta & Shulman, 2002; Corbetta, Patel & Shulman, 2008). While these two networks are anatomically distinct, the main nodes of both networks are located in the frontal and parietal association cortices.

A variant of this frontoparietal attention framework derives from non-invasive EEG recordings in humans (Li et al., 2010) and from single-neuron recordings in non-human primates (Buschman & Miller, 2007). This work focuses on the respective roles of the frontal and parietal association cortices in Search versus Pop-out. The central prediction that follows from these studies is that Search preferentially engages frontal cortex whereas Pop-out preferentially engages parietal cortex.

A challenge to a pure frontoparietal model of visual search has recently been introduced in a series of publications in non-human primates. These studies demonstrate that visual search of natural scenes rely extensively on medial temporal lobe (MTL) structures such as the entorhinal cortex (Killian et al., 2012; 2015). According to Killian and colleagues, visual search belongs to the family of *navigation* behaviors, not merely visual attention behaviors. In other words, visual search in primates can be conceptualized as navigation in visual space, analogous to navigation in physical space in rodents (Meister & Buffalo, 2016, Nau et al., 2018). Consistent with this, a recent study examining single-neuron responses in the human hippocampus and amygdala reported that individual neurons in these regions reflect target detection processes during visual search of natural images (Wang et al., 2018).

A second reason to predict MTL engagement in visual search, and especially in the visual pop-out phenomenon, stems from its association with novelty detection. Hedwig von Restorff discovered the relationship between novelty and memory nearly 90 years ago (von Restorff, 1933). Subsequent findings established that MTL memory structures are necessary for the detection and recollection of novel items (Knight, 1996; Parker et al., 1998). These findings raise the question of whether classical, well-parameterized experiments targeting Search and Pop-out also engage MTL structures.

A study of four participants with brain lesions that included the MTL (Chun & Phelps, 1999) showed results consistent with this prediction: Patients with MTL lesions were not able to benefit from repetition of search arrays (where all distractors were identical from block to block), suggesting MTL involvement in implicit memory for visual spatial context in search. However, direct electrophysiological evidence for MTL engagement in visual search and pop-out in humans is lacking. This may be due in part to the neglect of the contributions of these structures to attention in the cognitive neuroscience literature. In addition, the reduced signal-to-noise ratio in the MTL in fMRI recordings in humans impairs researchers’ ability to draw conclusions about putative MTL involvement in various cognitive tasks. In this study, we address the question of MTL involvement in Search and Pop-out using direct intracranial recordings in humans.

### Do Search and Pop-out map differentially onto the frontal and parietal cortices, respectively?

Returning to the question of Search and Pop-out as two separable perceptual processes, the idea that humans possess two visual systems goes back at least four decades (Treisman & Gelade, 1980; Julesz, 1981). These two putative perceptual systems have been proposed to map onto two distinct neural systems. Several models of the anatomical substrates for these two putative systems exist: and we discuss three of these models below.

First, a set of findings from non-invasive EEG in humans, and electrophysiology in primates, suggest that Pop-out is implemented in parietal cortex (Li et al., 2010; Buschman & Miller, 2007, but see Nobre et al., 2002). This is consistent with reported salience maps in lateral intraparietal cortex (area LIP) in monkeys (Gottlieb et al., 1998); and a broader role for parietal cortex in detecting external visual stimuli, holding such information in memory, and generating motor action plans (Mangano et al., 2015; Xu, 2018; Regev et al., 2018; Martin et al., 2019). By contrast, prefrontal cortex (PFC), including the frontal eye fields (FEF) and lateral frontal cortex are implicated in Search (Leonards et al., 2000; Buschman & Miller, 2007; Rossi et al., 2007; Li et al., 2010). Consistent with this view, these areas have also been shown to contain topographic visual maps (Kastner et al., 2007; Silver & Kastner, 2009; Mackey et al., 2017).

Second, Corbetta’s and Shulman’s (2002; Corbetta et al., 2008) model of visual attention, focuses on *networks* for goal-directed versus stimulus-driven attention, incorporating subregions of several cortical lobes. They propose that the DAN, which includes the intraparietal sulcus (IPs) and the FEF, supports goal-directed attention as deployed in Search. According to Corbetta’s and Shulman’s framework, the VAN - including the temporoparietal junction (TPJ), parts of the middle frontal gyrus (MFG), inferior frontal gyrus (IFG), frontal operculum, and anterior insula - are involved in stimulus-driven attention as deployed in Pop-out.

Third, at least two theoretical models exist for the implementation of Pop-out. One model proposes that the central mechanism enabling the visual pop-out phenomenon is computation of salience in early visual cortex, most notably V1 (Zhaoping, 2002; Zhaoping & Dayan, 2006; Zhaoping, 2019). An opposing model posits that Pop-out is, instead, a top-down phenomenon. This gives rise to the prediction that there should be neural correlates of Pop-out in higher-order cortical areas such as frontal cortex (Hochstein & Ahissar, 2002), consistent with the observation that PFC lesions cause decreased EEG novelty responses in humans (Knight, 1984).

As a fourth perspective, recent empirical evidence emphasizes the similarities rather than the differences between the tasks. Both fMRI work (Leonards et al., 2000) and a single intracranial study (Ossandon et al., 2012) suggest that the networks supporting Search and Pop-out are remarkably similar, and centered on fronto-parietal regions including the DAN. Local neural differences including greater activation in lateral frontal cortex in Search were observed (Leonards et al., 2000), perhaps reflecting increased working memory demands. The fact remains, however, that Search versus Pop-out engender robust behavioral differences, and the origin of these differences must reside somewhere in the brain.

In the present study, we employ direct intracranial recordings of neural activity to map the anatomical sites at which Search and Pop-out converge and diverge in the human brain. Intracranial recording provides a method with improved spatiotemporal resolution compared to non-invasive neural recording methods in humans (Parvizi & Kastner, 2018). This superior temporal resolution enables us to observe and reject artifacts driven by RT differences across attention conditions, which may have confounded the results in previous studies using lower-resolution recording methods. We analyzed intracranially recorded voltage signals from 1,321 ECoG and SEEG electrodes in 23 patients with medically refractory epilepsy, each of whom were undergoing diagnostic recording in preparation for potential resective surgery. Across patients, the sensors captured extensive areas of cortex as well as the MTL and the amygdala. This combination of high spatiotemporal resolution and broad coverage positions this study to assess where and how these two modes of searching our visual environment differ. We ask which regions of the brain differ for Search and Pop-out and assess to what degree these differences overlap with existing models of the implementation of Search and Pop-out in the brain.

The patients completed a simple visual search task (Figure 1), with two experimental conditions, Search and Pop-out. We focused on task-related activity increases, and condition-related modulations, in high-frequency activity (HFA; 80-150 Hz). HFA has been shown to correlate both with the fMRI BOLD signal (Logothetis et al., 2001; Mukamel et al., 2005; Nir et al., 2007) and with multi-neuron activity (Ray et al., 2008; Ray & Maunsell, 2011). The findings demonstrate a robust role for the MTL both in Search and Pop-out, and highlight sub-regional differences within the PFC, including different activation profiles between superior and inferior lateral PFC.

**Figure 1.**
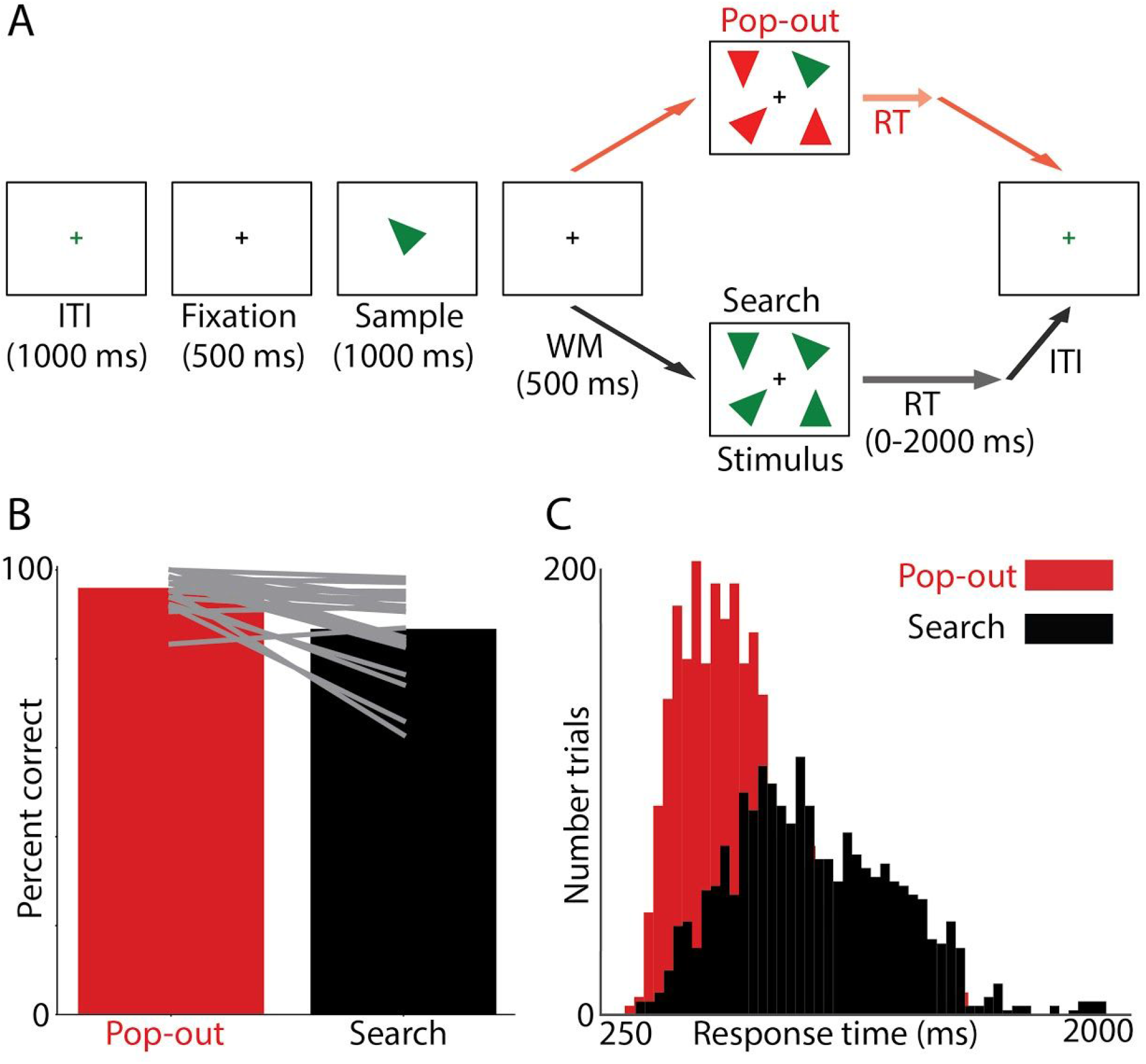
A) Visual search task adapted from Li et al. (2010). Patients searched for a target triangle (‘Sample’) of a given color and orientation. Upon locating the target triangle, they indicated its location on the left or right half of the screen by button press. ITI, Inter-Trial Interval; Fixation, Fixation Interval; Sample, Sample Interval; WM, Working Memory Interval; Stimulus, Stimulus Display Interval; RT, Response Time. B) Proportion correct responses in the Pop-out (red) and Search (black) condition across the 23 patients included in the study. Grey lines indicate individual subjects’ performance. C) RT histograms from all correct trials for all participants (N=23), showing the Pop-out (red) and Search (black) conditions.

## Method

### Participants

Thirty participants were enrolled in the study while undergoing treatment for medically refractory epilepsy at Oslo University Hospital, Norway (N = 8), California Pacific Medical Center, USA (N = 7), or UC Irvine Medical Center, USA (N = 15). The Institutional Review Boards at each hospital as well as the Committee for the Protection of Human Subjects at the University of California, Berkeley, approved the study procedures. In Norway the study was also approved by the Regional Committees for Medical and Health Research Ethics. Each patient provided written informed consent prior to participation. Electrode placement was determined solely by clinical needs.

Seven patients’ datasets were excluded from the analyses. Two were excluded due to chance behavioral performance in the experiment. One was excluded for having fewer than 15 trials in one of the two experimental conditions, after applying trial exclusion criteria (see section *Removal of epochs* below). Two patients were excluded for showing no task-related activity beyond primary sensory areas in the recordings (see section *Task-active electrode selection: Identifying significant increases*), and two patients were excluded for excessive high-frequency noise.

Of the remaining 23 participants (35.04 ± 13.97 years; mean ± SD; 13 female), 21 were right-handed, and 2 were ambidextrous (see Supplementary Table 1). For the patients who participated in the U.S. (n = 17), twelve were native English speakers; two were native Spanish speakers; one was bilingual in English and Spanish; one was an ASL speaker; and one was a native Vietnamese speaker. All patients whose data was collected at Oslo University Hospital (n = 6) were native Norwegian speakers. Task instructions were provided in the participant’s first language, either by the experimenters or with the help of a family member. Language was not a barrier to task comprehension in any of the subjects. All participants had IQ > 85; normal or corrected to normal vision; and no known deficits in visual perception or color vision.

### Stimuli

Participants completed a visual search task adapted from Li et al. (2010; 2013). The stimuli consisted of acute, isosceles triangles that were either red or green, and presented on a white background (Figure 1) on a Windows laptop (15.6″ LCD screen). The stimuli were presented, and responses recorded, using E-Prime 2.0 software (Psychology Software Tools, Sharpsburg, PA).

The laptop was placed in front of participants at a viewing distance of 40-60 cm (approximately 16-24″). Each triangle had a base of 4.0 cm (3.8-5.7° visual angle) and a height of 4.5 cm (4.3-6.4° visual angle). On the stimulus display, the distance of each triangle from a central fixation cross was 6.0 cm (5.7-8.6° visual angle) horizontally and 4.0 cm vertically (3.8-5.7° visual angle), measured between the fixation cross and the center of each stimulus triangle.

To synchronize the neural and behavioral recordings for subsequent analyses, analog channels transmitted stimulus onset and offset signals from the stimulus computer to the neural recording hardware. At the hospitals in the U.S., a photodiode was used to detect light changes on the stimulus monitor, and at Oslo University Hospital, a TRS (“Tip, Ring, and Sleeve”, see: https://missionengineering.com/what-is-a-trs-cable/) connector transmitted an audio signal from the stimulus computer to the recording rig.

Participants were instructed to locate a target triangle among four candidate triangles on the screen. The target triangle was defined by a color (red or green) and an orientation (1 of 8 orientations: 0°, 45°, 90°, 135°, 180°, 225°, 270°, or 315°). Patients indicated whether they found the target triangle on the left or right half of the screen by pressing the left versus right arrow key on the laptop keyboard, or the left versus right button of an external mouse, depending on the physical constraints of the recording room.

The trial sequence started with a green fixation cross (1000 ms, *Inter-Trial Interval*; notice that a 500-ms *Baseline Interval* was a subset of this interval), followed by the fixation cross turning black (500 ms, *Fixation Interval*). Next, the target triangle, which participants were required to hold in memory, was presented in the center of the screen (1000 ms, *Sample Interval*), followed by a fixation cross (500 ms, *Working Memory Interval;* Eimer, 2014), and ultimately the stimulus display consisting of four triangles including the target triangle (*Stimulus Display Interval*). The trial was terminated after the response or, if the participant did not respond, after 2,000 ms (Figure 2.A).

**Figure 2.**
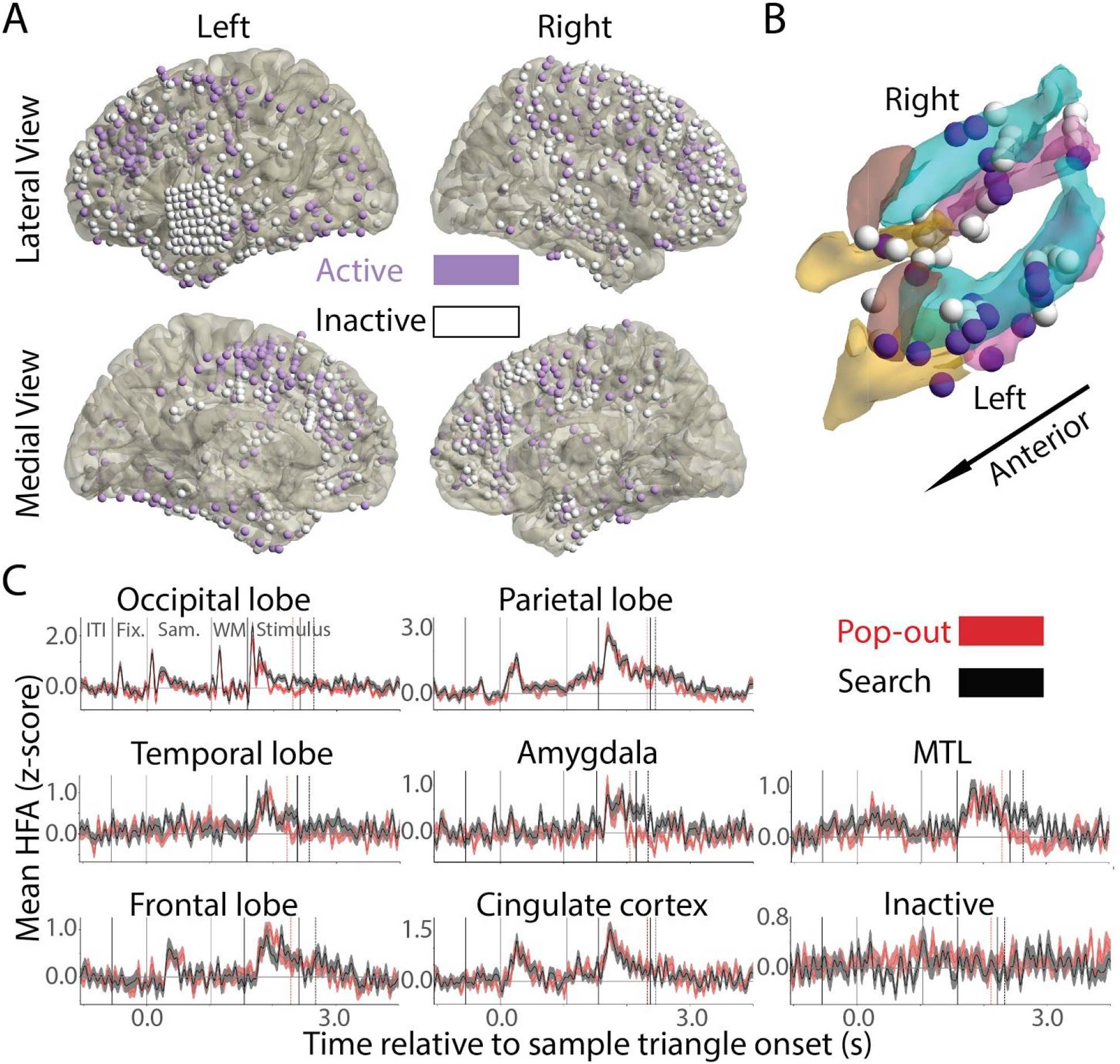
Task-selective electrode sites across the dataset (N=363 of 1,160). A) Task-active (purple) and inactive (white) sites visualized on a common brain reconstruction (Colin-27). B) Task-active electrodes in the medial temporal lobe. Active electrodes are plotted in purple, and inactive electrodes are plotted in white. C) Example time courses from seven major regions of interest, and an example time course from an inactive electrode site. Pop-out time courses are plotted in red and Search time courses in black. The vertical red dashed line indicates the median RT in the Pop-out condition. The vertical black dashed line indicates the median RT in the Search condition. The various task epochs are indicated on the top left panel, and correspond to those shown in Figure 1.A: ITI, Inter-Trial Interval; Fix., Fixation Interval; Sam., Sample Interval; WM, Working Memory Interval; Stimulus, Stimulus Display Interval. Note that purple was used (rather than red and green) to improve contrast for colorblind readers.

Each experiment included a total of 128 trials, divided into four blocks of 32 trials each. The task included two experimental conditions, Search and Pop-out. In Search, all four triangles had the same color (all red or all green), while in Pop-out, the target triangle had a different color than the three distractor triangles (red target with three green distractors, or green target with three red distractors). Half the trials (n = 64) were Search trials. They were randomly interspersed with Pop-out trials, so that participants could not anticipate which condition an upcoming trial belonged to. Trial order was randomized separately for each participant.

### Behavior analysis

The behavioral outcome measures were response accuracy (proportion correct) and response time (RT, *ms*). To compare behavioral performance between the two conditions, we computed a paired *t*-statistic between percent accuracy in the Pop-Out versus Search conditions, excluding non-response (time-out, >2000 ms from stimulus onset) trials. We assessed significance using a permutation test, where we shuffled condition labels 100,000 times, and then we compared the *t-*statistics obtained in the permutation tests to the observed *t*-statistics. The *p*-value was computed as: (1 + number of permuted statistics more extreme than the observed statistic) / (1 + number of permutations). We calculated the same test to compare median RTs between the two conditions across subjects, including only correct-response trials.

### Neural data acquisition

Intracranial electrodes (ECoG or SEEG) were implanted for approximately one week in each patient to determine the epileptogenic focus. In eight patients, ECoG arrays, organized either in two-dimensional grids or one-dimensional strips, were placed on the cortical surface. In fifteen patients, SEEG depth electrodes, targeting subcortical structures, were implanted. A total of 1,321 electrodes were analyzed across the 23 participants.

At Oslo University Hospital, ECoG and SEEG data were recorded using two 64-channel NicoletOne (Natus Neuro Inc., Middleton, WI, USA) amplifiers (in four patients) and a 256-channel ATLAS (Neuralynx, Bozeman, MT, USA) digital acquisition system (in one patient). For the SEEG cases (n = 5), the exposed electrode diameter was 0.8 mm, and the center-to-center distance between electrodes was 3.5 mm. In four of these patients, the digitization rate was 512 Hz (NicoletOne system). In one patient, it was 16,000 Hz (ATLAS system). ECoG data were sampled at 1,024 Hz (NicoletOne system). The electrodes were manufactured by DIXI Medical (Besançon, France).

At CPMC, ECoG data were recorded using a Nihon Kohden (Tokyo, Japan) Neurofax EEG-1200 digital acquisition system with 128/256-channel amplifier capacity. The digitization rate was 1,000 Hz. All five patients at this site were implanted with surface contacts (grids and strips). The electrodes were manufactured by the Ad-Tech Medical Instrument Corporation (Oak Creek, WI, USA).

At UC Irvine Medical Center, data were recorded using a Nihon Kohden digital acquisition system with a 128/256-channel amplifier at a digitization rate of 1,000/5,000 Hz, for both ECoG and SEEG cases. For the ten patients implanted with SEEG depth electrodes, the diameter of the electrode contacts was 0.9 mm, and the inter-electrode distance was 5.0 mm. The electrodes were manufactured by Integra Life Sciences (Plainsboro, NJ, USA).

For all patients recorded with surface electrodes, the exposed electrode diameter was 2.3 mm. The inter-electrode spacing was 10.0 mm for all ECoG grids in all patients, except a single 8×8 high-density grid in one patient, where the center-to-center spacing was 4 mm.

### Anatomical reconstruction and visualization

For electrode localization, we first segmented each patient’s preoperative T1-weighted MRI scan using Freesurfer 5.3.0 (Dale et al., 1999). Next, we fused the MRI image with a post-implantation CT scan using the Fieldtrip toolbox (Oostenveld et al., 2011; Stolk et al., 2018). To correct for the displacement of electrodes and brain tissue due to pressure changes related to the patient’s craniotomy, electrodes were realigned to the preoperative cortical surface (Hermes et al., 2010; Dykstra et al., 2012). We inspected the individual fusions for maximal interlocking between the CT and MR visually for quality assurance. In the case of two patients, whose native-space MRIs were missing, we co-registered the post-implantation CT scans to the stereotaxic Colin-27 brain template (Collins et al., 1998).

For group-level visualization, surface-based normalizations were conducted using Freesurfer via cortical gyrification patterns, and volume-based normalizations through fusion to the Colin-27 brain via overall geometry patterns. By doing so, we linked patient brains to their template homologs, allowing accurate comparison of regions of interest (ROIs). In SEEG datasets, for which each physiological time-series was calculated using a bipolar montage between pairs of electrodes, we calculated the position of each electrode as the mid-point between the two original electrode locations in native space (Burke et al., 2013; Jafarpour et al., 2019). To visualize electrodes showing task-related effects, we used FieldTrip and custom Matlab scripts.

### Assignment of electrode locations to anatomically defined regions of interest

A neurologist (RTK) identified the locations of the individual electrodes based on inspection of the fused MR and CT images, displayed in the patient’s native space, using the BioImage Suite toolbox.

We identified electrodes as belonging to one of the following eight broad ROIs (see Supplementary Table 2): Frontal cortex (not including sensorimotor cortex), parietal cortex (not including sensorimotor cortex), sensorimotor cortex, temporal cortex, occipital cortex, cingulate cortex, medial temporal lobe (MTL), and amygdala. Sensorimotor cortex, which was engaged due to the behavioral response (a button press), was not of primary interest in this study, and was not further examined. We additionally examined activations in subregions of those ROIs that had greater than 50 electrodes (Supplementary Table 2).

In frontal cortex (Supplementary Table 3), we examined the following five subregions, defined by gyral and sulcal landmarks: (1) the superior frontal gyrus and sulcus, (2) the middle frontal gyrus, (3) the inferior frontal gyrus and sulcus, (4) the orbitofrontal cortex, and (5) the medial prefrontal cortex.

In parietal cortex (Supplementary Table 4), we separately examined activations in: (1) the superior parietal lobule including the intraparietal sulcus, (2) the inferior parietal lobule comprised of the supramarginal gyrus, angular gyrus, and temporoparietal junction, and (3) the precuneus.

In temporal cortex (Supplementary Table 5), we examined the following seven subregions: (1) the insula, (2) the superior temporal gyrus, (3) the superior temporal sulcus, (4) the middle temporal gyrus, (5) the inferior temporal gyrus including the middle temporal sulcus, (6) the ventral stream including the lingual and fusiform gyri, and (7) the temporal pole.

In cingulate cortex (Supplementary Table 6), we examined (1) the anterior cingulate cortex, (2) the midcingulate cortex, and (3) the posterior cingulate cortex.

Finally, in the medial temporal lobe (Supplementary Table 7), we examined the (1) hippocampal formation (HF) including the hippocampus and subiculum, (2) parahippocampal cortex, and (3) entorhinal cortex including perirhinal cortex.

### Preprocessing of neural data

We recorded the local field potential from a total of 503 ECoG and 818 SEEG electrodes in the 23 participants included in the final study sample. In subjects with an original sampling rate of 5,000 or 10,000 Hz, we resampled the signal to 1,000 Hz. The resulting datasets had sampling rates of 512, 1,000, or 1,024 Hz. To remove slow drift and high-frequency noise respectively, we high-pass filtered the signal at 1 Hz, and low-pass filtered it at 180 Hz. To remove line noise, we notch-filtered the signal at 60 Hz and harmonics for datasets recorded in the U.S., and at 50 Hz and harmonics for datasets recorded in Norway.

Each recording was visually inspected by a neurologist (RTK) for epileptic activity or poor signal quality (such as detached electrodes or high-frequency noise). Electrodes that reflected signal from epileptic tissue or white matter, or were consistently noisy during the recording, were removed from the dataset. Temporal epochs that showed epileptic activity spread were removed across all electrodes. To remove remaining shared noise sources from the data in the surface (ECoG) datasets, we applied the common average reference (subtracting the point-by-point average signal of the preprocessed dataset from each time point of retained electrodes) to all grids within patient. In depth (SEEG) datasets, we applied a bipolar reference (pairwise subtraction of adjacent electrode time-series).

### Spectral decomposition

We extracted the analytic amplitude of the HFA signal through three steps. First, we bandpass-filtered the time-series from the complete recording in each electrode between 80 and 150 Hz using a zero-phase FIR filter (mne.io.Raw.filter function from the MNE toolbox, https://martinos.org/mne/stable/generated/mne.io.Raw.html). We then computed the Hilbert transform of the filtered signal (mne.io.Raw.apply_hilbert function from the MNE toolbox) yielding a complex time-series, of which we took the absolute value to obtain the instantaneous analytic amplitude. Finally, we low-pass filtered the HFA analytic amplitude at 10 Hz to facilitate detection of temporal variation on a time-scale of approximately 100 ms, following Haller et al. (2018).

### Baseline normalization

To examine the relationship between the behavioral and neural data, we segmented the full recording time-series into trial epochs, time-locked to the onset of the sample triangle (start of the *Sample* interval). A full trial epoch ranged from 1,050 ms before the onset of the sample triangle to 4,000 ms after. We defined the *Baseline Interval* as the first 500 ms of the trial, i.e., -1,050 to - 550 ms relative to the onset of the sample triangle. During the baseline, the participant watched a black fixation cross at the center of the monitor. We ended the trial epoch at 4,000 ms, because this captures the longest trials plus a 500-ms period of response and post-response activity. To facilitate detection of task-related changes in neural activity, we normalized the HFA analytic amplitude to the neutral *Baseline Interval*, by computing an adapted z-score as follows: From each sample point of the full trial epoch, we subtracted the mean, and divided by the standard deviation, of the pooled baseline interval (across all included baseline epochs within condition and within each electrode). We then separated trials belonging to the Pop-Out versus Search condition, and conducted all subsequent analyses separately within each condition.

### Removal of epochs

We excluded epochs from the dataset according to the following criteria: incorrect behavioral response, no response prior to trial timeout, epileptic activity or consistent noise in the raw time-series, RT outliers, and HFA activity outliers. RT outliers were defined as trials where the RT was more than 3 interquartile ranges (IQR, i.e., the 75th percentile - 25th percentile) lower than the 25th percentile or more than 3 IQR higher than the 75th percentile of the subject’s own RT.

In order to minimize the number of trials with HFA signal artifacts in electrodes of interest, while maximizing the total number of trials retained across a patient’s dataset, we used a two-step approach for identifying HFA activity outliers. First, we computed the set of active electrodes (see section, *Task-active electrode selection: Identifying significant increases* and Supplementary Figure 1) in each patient’s dataset prior to HFA outlier exclusion. Next, we excluded trials that showed a max-min range greater than 6 IQR above the 75th percentile (or greater than 6 IQR below the 25th percentile) in any one active electrode. Trials were excluded across each patient’s full dataset. The maximum number of trials that were excluded in any one patient based on this criterion was 8 across the dataset, and 4 in any one condition. The smallest number of trials retained in any one subject and condition after all trial exclusion criteria was 21. Finally, we computed the set of active electrodes a second time, on the data where HFA outliers had been removed. The max-min range was selected as a metric of deviation in order to capture the time course shape of the most common HFA signal artifacts (rapid, high-amplitude modulations). The threshold of 6 IQR was chosen based on examination of two patients’ datasets, and then generalized across all patients.

### Task-active electrode selection: Identifying significant increases

We used a permutation test approach adapted from Maris & Oostenveld (2007; Supplementary Figure 1) to select the subset of electrodes that showed task-relevant changes in HFA activity for further analyses. We considered an electrode that showed a significant task-related increase relative to baseline in either condition to be task-active. We also identified electrodes that showed task-related decreases using an equivalent procedure, but these effects are not the main focus of the current paper. To make the comparison, we selected two intervals (“task interval” below) of equal length to the baseline: (1) stimulus onset to 500 ms following stimulus onset, (2) 500 ms before response up to response. The significance calculation for each electrode was computed as follows: (1) Baseline-normalize both baseline and task intervals as outlined in the section, *Baseline normalization*, above. (2) For both the baseline and task intervals, compute a one-sample *t*-test across trials at each time point. (3) Define a cluster as the set of *t*-statistics associated with any set of two or more consecutive significant time points (p < 0.05; see Supplementary Figure 1, grey shaded area in the center panels depicting *t*-statistics). Select all clusters. (4) Compute the sum of *t*-values in each cluster. (5) Select the cluster with the largest sum of *t*-values (in both the baseline and the task intervals). (6) Subtract the largest sum of *t*-values in the baseline interval from the largest sum of *t*-values in the task interval, to obtain an observed statistic for the electrode. (7) Randomly permute the task versus baseline labels 1000 times, and repeat the calculations in steps 1-6 above. (8) Compute a *p*-value for each electrode as: (1 + number of permuted statistics larger than the observed statistic) / (1 + number of permutations). (9) An electrode with a *p*-value < 0.025 one-tailed (FDR-corrected across all electrodes in all subjects) in either of the two intervals in either condition was considered significant. We selected a cutoff of *p* = 0.025 to Bonferroni-correct our threshold. We chose this cutoff because we computed increases and decreases as two separate, but parallel, one-tailed tests (see above). For physiological plausibility, we added a constraint that the duration of a cluster had to be, at a minimum, 50 ms.

### Significance-testing the anatomical distribution of task-selective effects

We tested the distribution of task-active electrode sites for significant regional or hemispheric differences, using chi-square tests, according to the following approach: First, we tested if there were any regional differences in the distribution of task-active effects among the four broadly defined cortical lobes, cingulate cortex, MTL, and amygdala (Supplementary Table 2). We similarly tested if there were differences between the subregions of the frontal cortex (Supplementary Table 3), parietal cortex (Supplementary Table 4), temporal cortex (Supplementary Table 5), cingulate cortex (Supplementary Table 6), and the MTL (Supplementary Table 7), again using chi-square tests corrected for multiple comparisons using the FDR-correction. We did not examine subregion effects for the occipital lobe or the amygdala, due to low electrode counts in these regions. For any omnibus chi-square test (Supplementary Tables 2-7) that was significant, we conducted follow-up pairwise chi-square tests to determine which ROIs or subregions were driving the overall effect. Any regions, for which the expected number of observations (electrodes) for either active or inactive status was less than 5, were excluded from the analysis. Specifically, the precuneus was excluded from the analysis of parietal subregions, and the temporal pole was excluded from the analysis of temporal subregions.

### Identification of electrodes showing condition-based effects

We separately identified electrodes that showed greater activity increases in Pop-out than Search (“Pop-out electrodes”) and electrodes that showed greater activity increases in Search than in Pop-out (“Search electrodes”). To accomplish this, we used a very similar procedure (Supplementary Figures 2 and 3) to the one employed for identifying task-active electrodes, except for the following modifications: (1) We considered only electrodes that showed a significant task-related increase (see *Task-active electrode selection: Identifying significant increases* above). (2) We excluded any electrodes located in sensorimotor cortex (localized according to the procedure described in the section, *Assignment of electrode locations to anatomically defined regions of interes*t) due to potential confounding effects of the behavioral response being given by button press. For example, it is conceivable that patients pressed the button with greater force in the easier and more confident Pop-out than Search condition, giving rise to spurious condition-related effects. (3) Because we were interested in which electrodes showed a significantly larger *increase* relative to baseline in one condition versus the other, we zero-clipped the signal prior to making the comparison (using the function numpy.clip()), so that all compared time points were greater than or equal to zero. Had we omitted this modification, we may have inadvertently identified electrodes as showing a pop-out effect when that effect was in fact driven by a temporary *decrease* in the Search condition rather than an *increase* in the Pop-out condition. (4) Instead of computing a one-sample *t*-test to identify significant deviations from 0 (step 2 in the task-active pipeline), we computed a two-sample *t*-test between the two conditions. (5) Instead of using two intervals as in the task-active pipeline, we considered a single interval: from stimulus onset to median RT across the two conditions. The reasons for this were that (i) we were no longer restricted to using a time interval of equal length to the baseline; (ii) we observed that most condition-related changes were of stimulus-related type; (iii) using the median RT as a cutoff seemed to be a reasonable choice in order to avoid undue bias in either direction by the selection of interval length.

Consistent with the task-active selection pipeline, we computed two one-tailed *t*-tests, one in each direction, at a *p*-value threshold of 0.025 for each direction. We FDR-corrected the *p*-values across all included electrodes (those showing a task-selective increase); and included a minimum duration cutoff of 50 ms, identically to the task-active pipeline above. Electrodes that showed a significant effect in both directions (n = 3) were not considered to show a condition-related effect, and were not further analyzed.

Different RT distributions between the two conditions are an integral feature of the task. However, specific artifactual condition differences in the neural data can occur as a result of these behavioral differences between the two conditions, as was also observed in a previous intracranial study on visual search (Ossandon et al., 2012). We removed these artifactual effects through the following steps: (1) We simulated three response types, which we predicted to occur in the HFA signal (previously reported in Ossandon et al., 2012; Haller et al., 2018): (i) stimulus-related responses; (ii) sustained responses; and (iii) response-related responses. We then tested how these response-types would behave given a representative patient’s RT distributions in the two conditions in this task. See Supplementary Figures 4-6. No artifacts driven by the known temporal characteristics of the ECoG/SEEG response (Ossandon et al., 2012; Haller et al., 2018, and simulated in Supplementary Figures 5-6) are expected in the case of stimulus-related responses. Sustained-type responses can give rise to artifactual Search effects. Response-related responses can give rise to artifactual Pop-out or Search effects. (2) We visually compared electrode traces that had been automatically labeled as showing a Pop-out or Search effect (using the pipeline described above) to the simulated artifactual responses. In cases where the signal time course of the selected traces matched a prototypical artifact time course, we labeled this effect as false. See Supplementary Figures 7-9 for example electrodes, for which the condition-effect labels were removed using this method.

## Results

### Behavior

Consistent with past studies using similar tasks (Buschman & Miller, 2007; Li et al., 2010; Ossandon et al., 2012), subjects were more accurate (*t*(22) = 4.73, p < 0.001) and faster (showed faster RTs, *t*(22) = -14.06, p < 0.001) in Pop-out than in Search (Figure 1). The mean accuracy in the Pop-out condition was 96.0 % with a standard deviation of 3.9 %. The mean accuracy in the Search condition was 86.8 % with a standard deviation of 9.6 %. The mean RT (of the median RTs from each patient) in Pop-out was 680.2 ms with a standard deviation of 146.5 ms. The mean RT in Search was 964.8 ms with a standard deviation of 152.2 ms.

### Task-selective responses by anatomically defined regions of interest

The total number of active sites, excluding electrodes implanted in sensorimotor cortex, was 363 out of 1,160, corresponding to 31% across the dataset (Supplementary Table 2). Task-selective responses were detected in all considered brain regions, including all four lobes of the neocortex; the cingulate cortex; the medial temporal lobe; and the amygdala (Figure 2).

The proportions of active electrodes differed significantly between these areas, χ2 (6, N = 1,160) = 41.84, *p* < .001 (Figures 2 and 3). The occipital lobe, with the most limited coverage (17 electrodes), showed the strongest activation profile (65%), as expected given the visual nature of the task. The proportion active electrodes in the occipital lobe was significantly larger than in the temporal lobe, χ2 (1, N = 386) = 14.70, *p* < .001 (FDR-corrected); and the cingulate cortex, (FDR-corrected). χ2 (1, N = 163) = 9.54, *p* = .008

**Figure 3.**
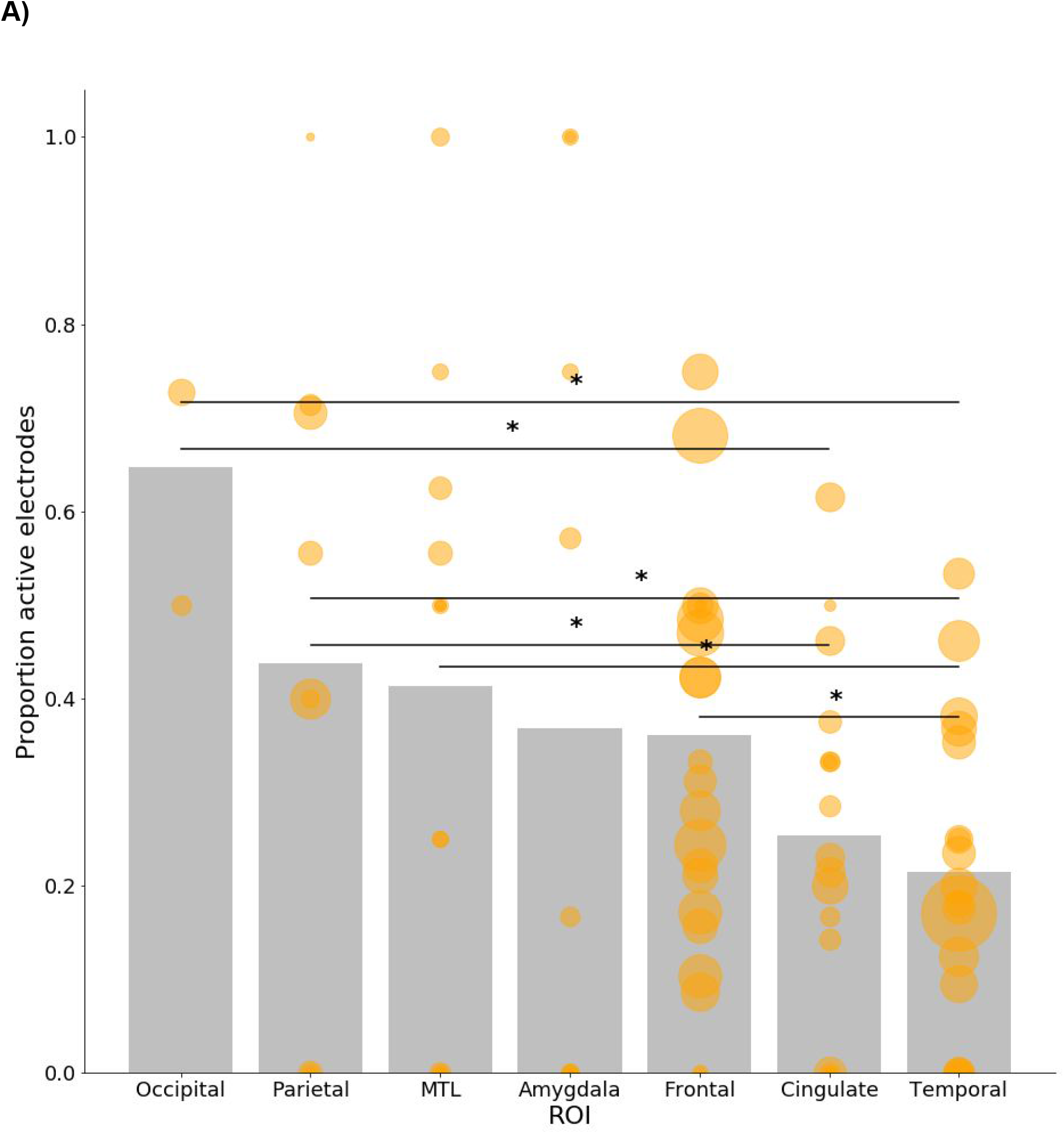
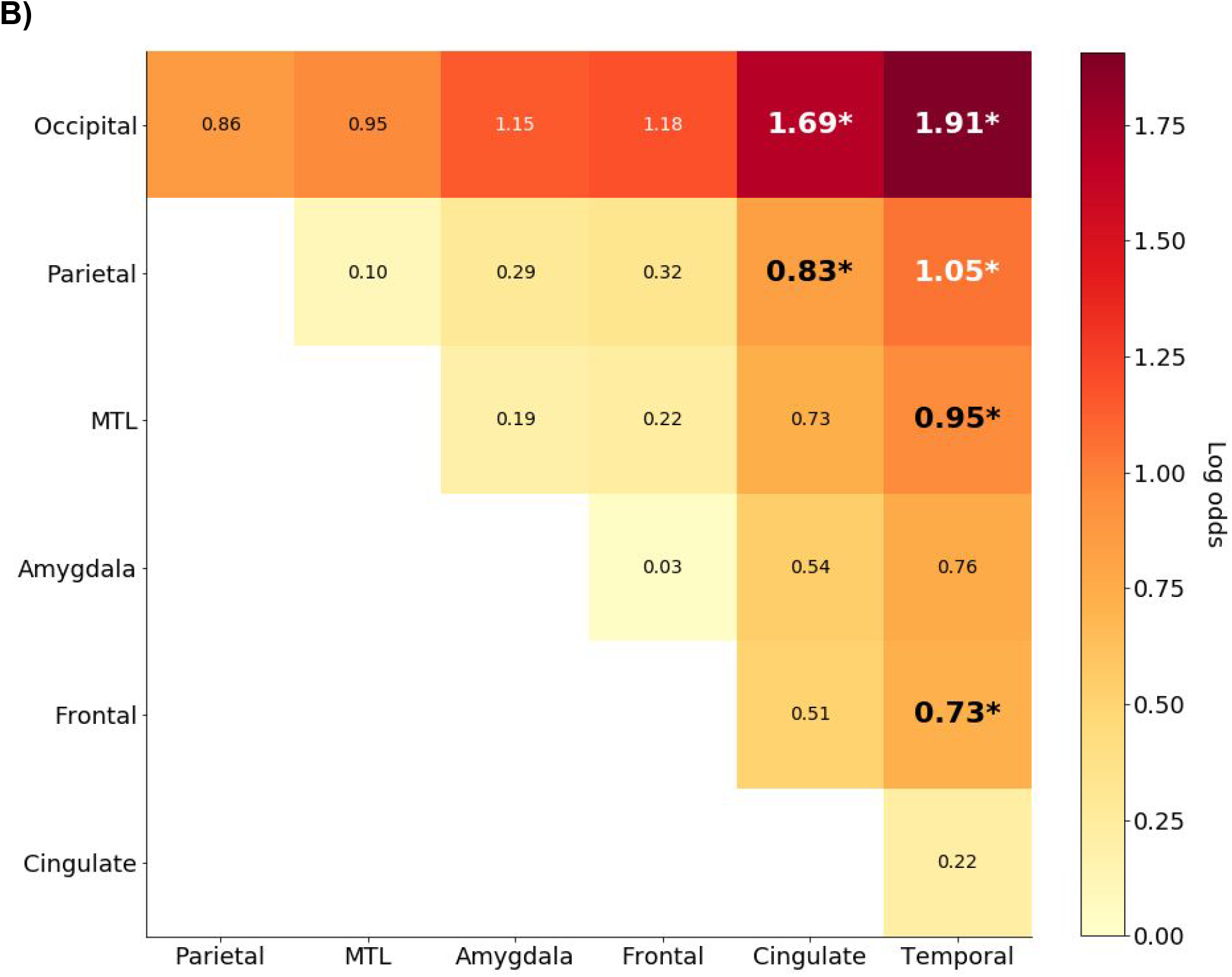
Distribution of task-active sites across the major ROIs. A) Proportion active electrodes by ROI. Bar height shows proportion active electrodes for a given ROI across the dataset. Circle height shows proportion active electrodes for individual subjects, who had coverage in that ROI. Circle area is proportional to the number of considered electrodes in that subject. Black asterisks indicate significant pairwise comparisons (*p* < 0.05). B) Effect sizes of pairwise comparisons between the major ROIs, corresponding to the pairwise chi-square test statistics reported. The effect sizes are illustrated as the log odds ratio between the ROI on the y-axis (indicating rows, on the left) versus the ROI on the x-axis (indicating columns, at the bottom): The proportion active electrodes in the ROI on the y-axis is always larger than the corresponding ROI on the x-axis. Statistics with an asterisk indicate a significant pairwise chi-square test.

Parietal cortex, extensively implicated in visual attention tasks, showed 44% active electrodes (Supplementary Tables 2 and 4; Figures 2 and 3). The proportion active electrodes in parietal cortex was larger than in the temporal lobe, χ2 (1, N = 449) = 16.16, *p* < .001 (FDR-corrected) and the cingulate cortex, χ2 (1, N = 226) = 7.24, *p* = .025 (FDR-corrected). We further examined three subregions of parietal cortex: the precuneus, the inferior parietal lobule (IPL), and the superior parietal lobule (SPL). In the precuneus, six of eight electrodes (75%) were active; due to the low electrode count, we did not further analyze this proportion. We compared the proportion active electrodes between the IPL (33% or 18/55 electrodes) and SPL (65% or 11/17 electrodes), and found a greater proportion of active electrodes in SPL, χ2 (1, N = 72) = 4.27, *p* = .039. The log odds ratio of this effect was 1.33. We note, however, that all SPL electrodes were derived from a single subject and the right hemisphere.

In PFC (Supplementary Tables 2 and 3; Figures 2 and 3), 36% of electrodes were active. This proportion was also larger than in the temporal lobe, χ2 (1, N = 821) = 20.28, *p* < .001 (FDR-corrected), but did not differ significantly from parietal sites. The proportion active electrodes were not uniformly distributed across frontal cortex, χ2 (4, N = 452) = 23.85, *p* < .001 (FDR-corrected; Figure 4). In lateral PFC, we observed a gradient of increasing activations from superior to inferior frontal gyri. In the superior frontal gyrus (SFG), we observed a proportional activation rate of 31%. In the middle frontal gyrus (MFG), 43% of electrodes were active. In the inferior frontal gyrus (IFG), the activation rate was 53%, with a larger proportion located in the left hemisphere (68% activation in the left hemisphere vs. 48% in the right hemisphere). The proportion active electrodes in IFG was larger than that in SFG, χ2 (1, N = 202) = 8.92, *p* = .009 (FDR-corrected). We observed proportionally fewer task-related increases in OFC (19%) in spite of extensive coverage (75 electrodes). This proportion was smaller than that in IFG, χ2 (1, N = 148) = 17.96, *p* < .001 (FDR-corrected), and MFG, χ2 (1, N = 211) = 11.32, *p* = .004 (FDR-corrected). In medial PFC (mPFC), 31% (12/39) of electrodes were active. This proportion did not differ significantly from other frontal subregions.

**Figure 4.**
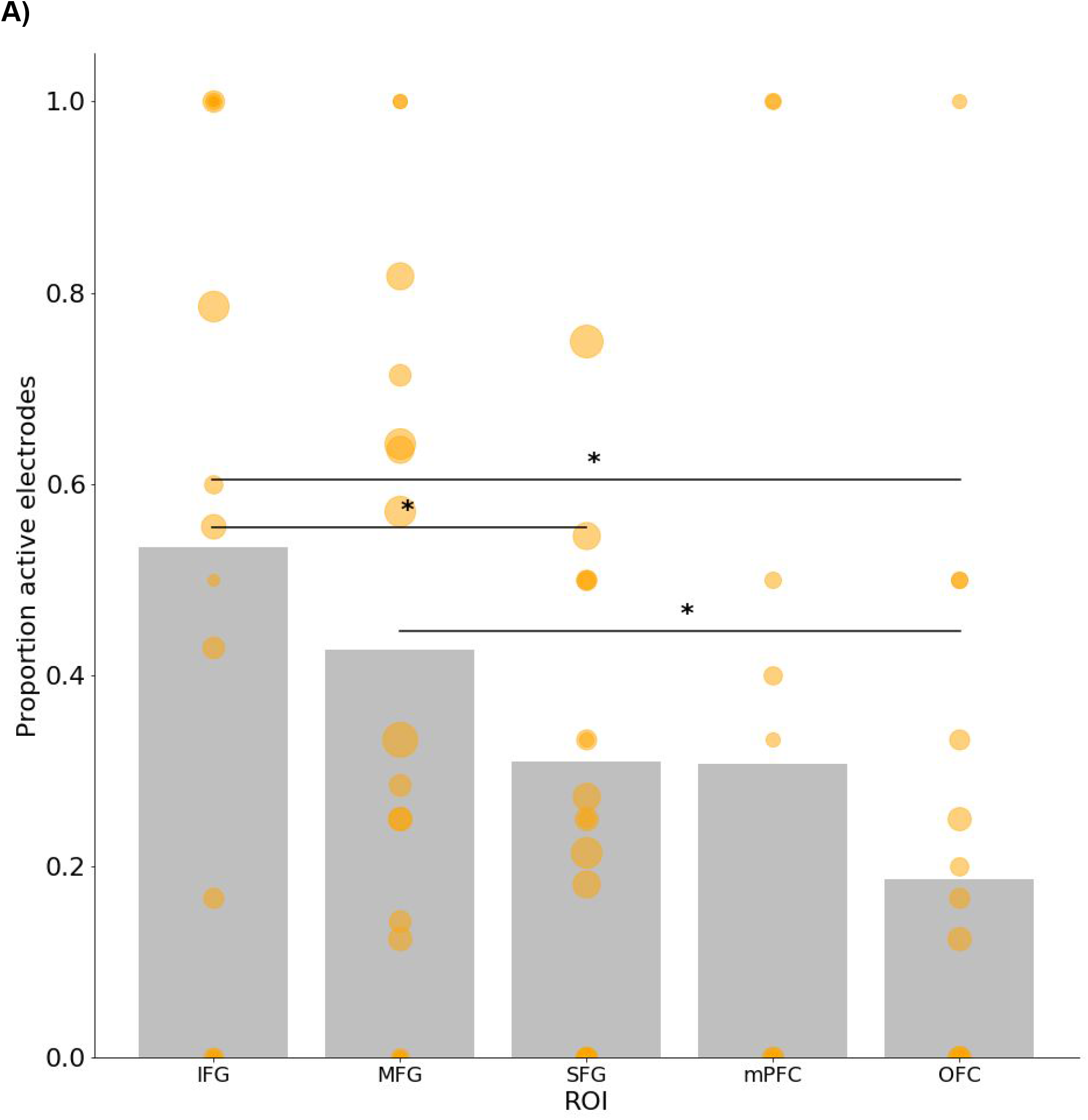
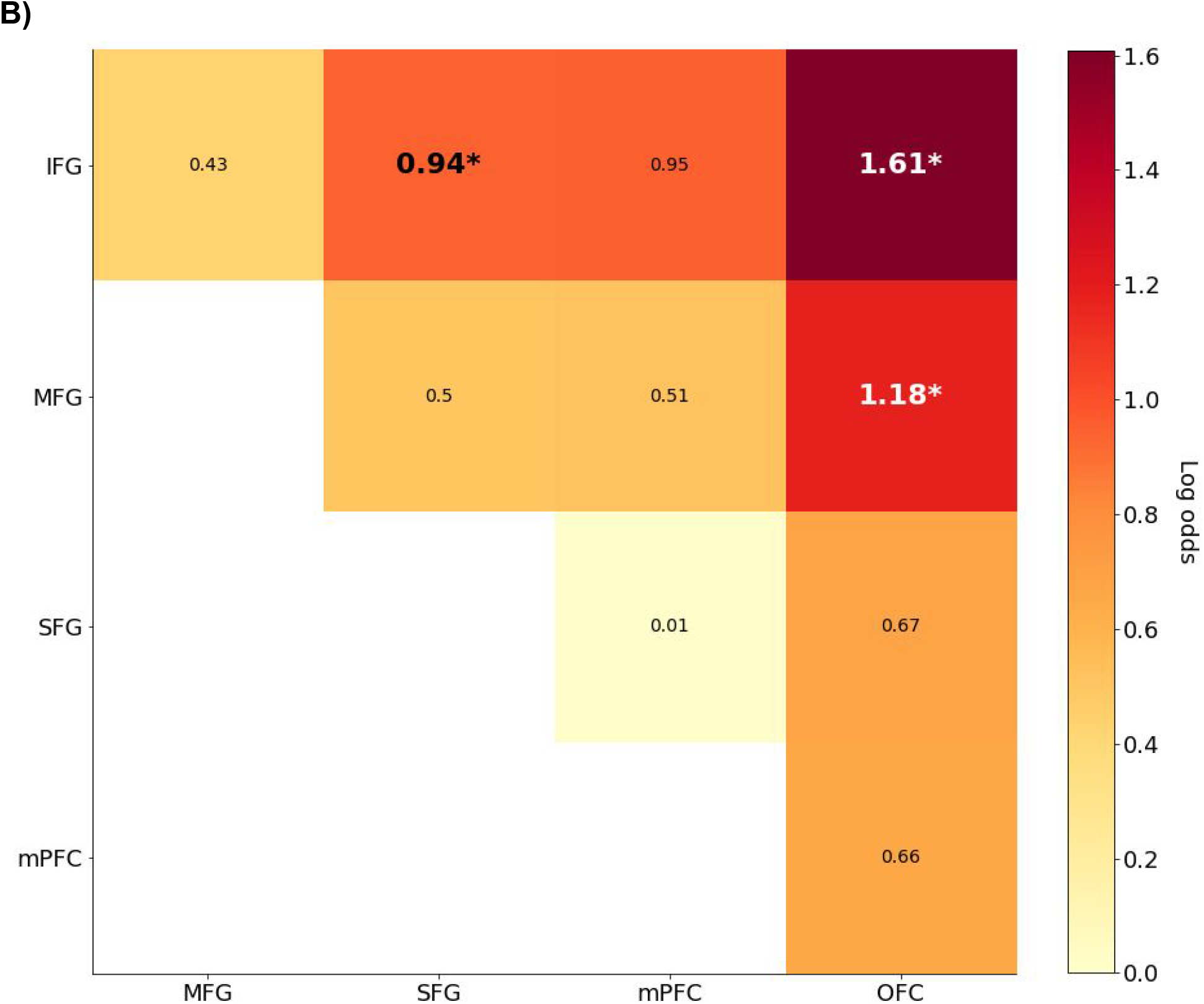
Distribution of task-active sites across subregions of frontal cortex. The layout of the figure corresponds to that of Figure 3. A) Proportion active electrodes by frontal subregion. B) Effect sizes of pairwise comparisons between frontal subregions.

The MTL (Supplementary Tables 2 and 7; Figures 2 and 3) showed a proportional activation profile at 41% (24/58 electrodes), indistinguishable from frontal cortex. Similarly to frontal cortex, this proportion was larger than that in temporal cortex, χ2 (1, N = 427) = 9.86, *p* = .008 (FDR-corrected). We detected no significant sub-regional differences within the MTL. In the hippocampus, we observed 41% (9/22) active electrodes. In parahippocampal cortex, 32% (7/22) of electrodes were active. Eight of fourteen (57%) entorhinal cortex electrodes were active. The amygdala (Supplementary Table 2, Figures 2 and 3) showed 37% active electrodes (14/38). This proportion did not differ significantly from the other ROIs.

The cingulate cortex (Supplementary Tables 2 and 6, Figures 2 and 3) was engaged only at 25% (37/146 electrodes). This proportion was smaller than that in the occipital and parietal cortices, as reported above. An omnibus chi-square test for significant regional differences between cingulate subregions was significant, χ2 (1, N = 146) = 7.12, *p* = .047 (FDR-corrected); however, the follow-up pairwise chi-square tests were not significant. We observed 33% (9/27) and 35% (18/52) active electrodes in the posterior (PCC) and midcingulate cortex (MCC), respectively. In the anterior cingulate cortex (ACC), we found only 15% active electrodes in spite of extensive coverage (10/67).

Temporal lobe engagement (Supplementary Tables 2 and 5; Figures 2, 3, and 5) was comparatively low, at 21% (79/369 electrodes). The proportion active electrodes in temporal cortex was reduced in comparison to occipital, frontal, and parietal cortices, as well as the MTL, as reported above. Within the temporal lobe, we observed a gradient of increasing activations from dorsal to ventral areas (Figure 5). This observation is consistent with the view that the ventral stream is central for object detection, while lateral temporal cortex would not be predicted to be engaged in visual search. The ventral stream areas (comprised of the lingual and fusiform gyri) showed 41% active electrodes (18/44). This proportion was significantly larger than that in STG, χ2 (1, N = 88) = 10.24, *p* = .010 (FDR-corrected), and STS χ2 (1, N = 126) = 13.51, *p* = .004 (FDR-corrected). The STG and STS showed only 9% and 11% active electrodes, respectively, in spite of extensive coverage in both areas (4/44 electrodes in STG and 9/82 electrodes in STS). The MTG showed 20% (16/79) active electrodes; ITG showed 22% active electrodes (13/59). In the insula, 26% (12/46) of electrodes were active. Of 15 electrodes localized to the temporal pole, 7 were active (47%). We did not consider the temporal pole in significance testing, due to low electrode count.

**Figure 5.**
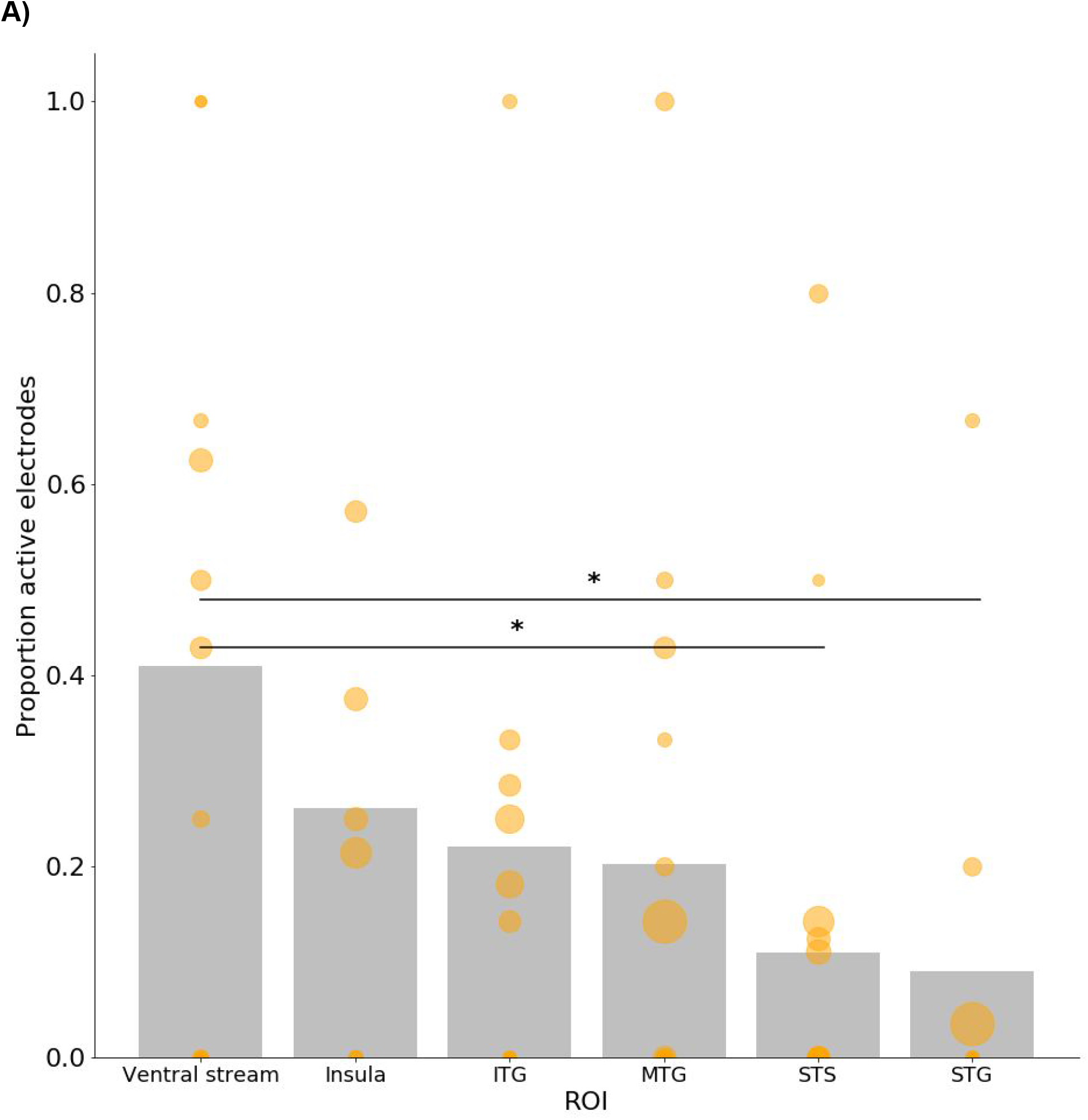
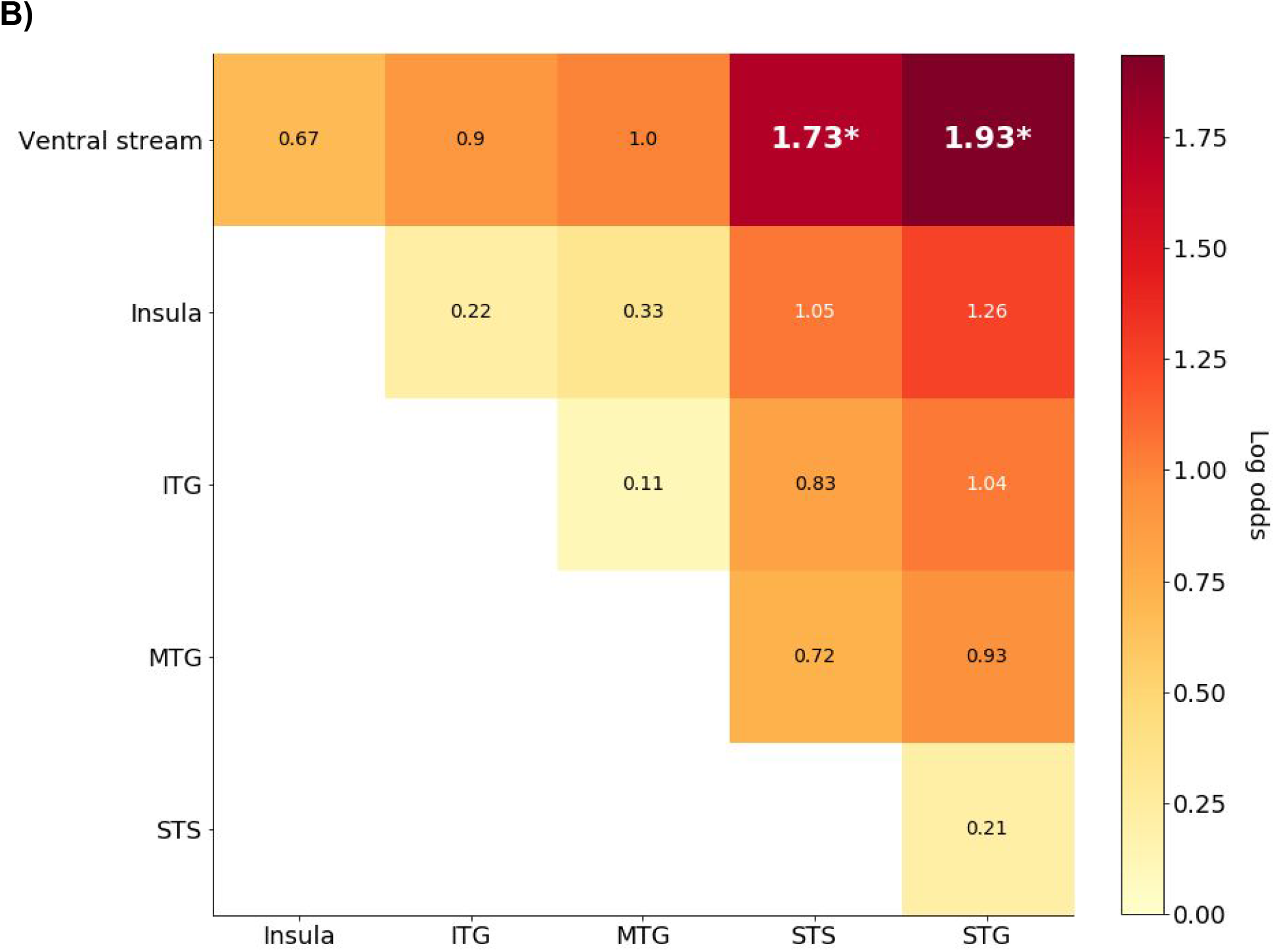
Distribution of task-active sites across subregions of temporal cortex. The layout of the figure corresponds to that of Figure 3. A) Proportion active electrodes by temporal subregion. B) Effect sizes of pairwise comparisons between temporal subregions.

In sum, occipital, frontal, parietal cortex showed an elevated proportion active electrodes compared to temporal cortex. The MTL showed a proportional activation rate on par with that in frontal and parietal cortex. As illustrated in Figures 3-5, none of these effects were driven by a single, anomalous subject.

### Condition-selective anatomical sites

Among the 363 electrodes which showed task-related increases outside of sensorimotor areas (Supplementary Table 2 and Figure 6), 30 sites (8.3%; 13 of 23 patients) showed condition-related modulations (*Pop-out > Search* or *Search > Pop-out*). This proportion is smaller than previously reported: Ossandon et al. (2012) found that 15% of electrodes with a sustained temporal profile in ROIs of the DAN showed condition differences, and 20% of electrodes with a stimulus-locked profile in the temporal and occipital lobes, showed condition modulations. This difference may be due to different methods for pre-selecting electrodes; determining statistical significance over time; and excluding condition modulations arising from RT-related artifacts. The total proportion of condition effects automatically detected in this study prior to artifact exclusion (65/363 electrodes or 17.9%) is similar to the proportions reported by Ossandon et al. (2012).

**Figure 6.**
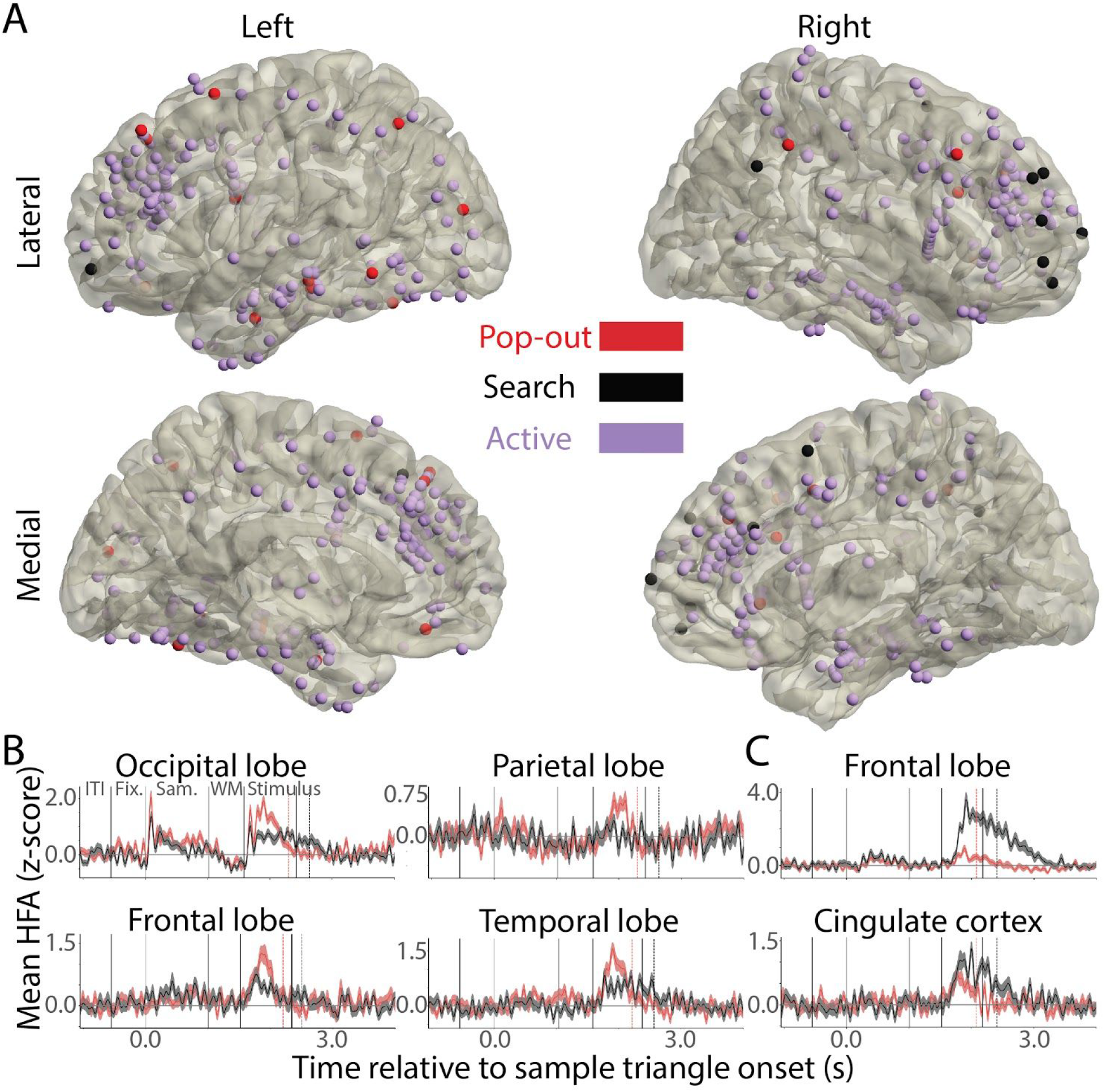
Condition-selective electrode sites. A) Pop-out (red) and Search-selective (black) sites are shown, along with task-active (purple) sites for which the two conditions did not differ. The brain template (Colin-27) is identical to that shown in Figure 2. B) Example time courses of electrode sites that showed a significant Pop-out selective effect. The mean signal in Pop-out is plotted in red, and Search in black. C) Example time courses of electrode sites that showed a significant Search-selective effect. The color scheme is the same as in (B).

Condition-related effects were found in all four cortical lobes as well as cingulate cortex. Because of the small number of sites showing condition-related modulations and their wide anatomical distribution, we do not statistically test these effects as a function of region, but rather report their observed anatomical locations directly.

Electrodes showing greater activity increases in the Pop-out than Search condition (n = 19 electrodes; 10 patients) were found in the parietal lobe (n = 4 electrodes; 4 patients; SPL, IPL, and SMG); frontal lobe (n=8 electrodes; 5 patients; SFG, superior frontal sulcus [SFS], MFG, IFG, OFC); temporal lobe (n=6 electrodes; 4 patients; MTG, ITG, insula, and fusiform gyrus), and occipital lobe (n=1 electrode; lateral occipital cortex). Electrodes showing greater activity increases in the Search than the Pop-out condition (n = 11 electrodes; 5 patients) were predominantly found in frontal cortex (9 electrodes; 4 patients). Of these frontal Search sites, all but one (mPFC) were found in lateral frontal cortex. The remaining two Search electrodes (N=2 patients) were in cingulate cortex (ACC and mid-cingulate cortex).

We removed specific artifactual condition differences, which occurred as a result of the different RTs distributions in the two conditions (see *Method* for details). Consistent with our simulations (Supplementary Figures 4-6), we observed three types of artifacts in the data: (1) spurious Search effects in electrodes with sustained activity (Supplementary Figure 7); (2) spurious Search effects in electrodes with response-related activity (Supplementary Figure 8); and (3) spurious Pop-out effects in electrodes with response-related activity (Supplementary Figure 9). We observed 14 electrodes showing the first type of artifact, located in all four cortical lobes, as well as the MTL and amygdala. Six electrodes showed the second type of artifact; these were located in frontal, occipital, and temporal cortex. Fifteen electrodes were characterized by the third type of artifact, and were found in frontal, parietal, and temporal cortex, as well as the cingulate cortex and MTL.

In sum, we found condition differences between Pop-out and Search at equivalent rates as those previously reported. Pop-out effects were found in all four cortical lobes, including frontal cortex. Search effects were found in lateral frontal cortex and cingulate cortex.

## Discussion

We used intracranial neural recordings with high spatiotemporal resolution in humans to document two previously undescribed properties of the visual search network in humans. First, there are marked regional differences in engagement of the PFC in this task, including an increasing gradient of activations from superior to inferior lateral prefrontal cortex. Second, the medial temporal lobe is engaged on par with the frontoparietal cortices in this classical attention task. In addition, we observe robustly Search-selective sites in lateral PFC, along with a distributed set of cortical sites selective for Pop-out. We discuss these findings below.

### Behavior results

Consistent with past research examining the neural correlates of serial search and pop-out, we observed faster RTs and higher accuracy in the Pop-out than the Search condition (Li et al., 2010; Ossandon et al., 2012; Yan et al., 2016). Moreover, the distributions of RTs in the two conditions were similar to those previously reported (Figure 1.C; Buschman & Miller, 2007; Li et al., 2010; Ossandon et al., 2012).

### Cortical distribution of task-active sites

#### Strong activation profile in visual areas

Across cortical areas, we observed the strongest activation profile in the occipital lobe (Figures 2 and 3) as would be expected in a visual experiment. Similarly, we observed strong activation in the ventral visual stream (Figure 5), consistent with its role in low-level vision, object detection, and fast visual categorization (DiCarlo & Cox, 2007; DiCarlo et al., 2012; Cauchoix et al., 2016).

#### Fronto-parietal network engagement

Both the frontal and parietal cortices were robustly active, as predicted by fronto-parietal models of attention allocation in both Pop-out and Search (Corbetta & Shulman, 2002; Buschman & Miller, 2007; Li et al., 2010; Ossandon et al., 2012). The robust parietal activations in the SPL and IPL (areas adjacent to the IPs) were also expected (Gottlieb et al., 1998; Silver et al., 2005; Silver & Kastner, 2009; Li et al., 2010).

#### Regional differences within prefrontal cortex

The lateral convexity of the human PFC is known to be involved in working memory and controlled attention (Knight et al., 1995; Barcelo et al., 2000; Curtis & D’Esposito, 2003). Lateral PFC is required for holding representations of sensory stimuli in mind over short periods of time (seconds) while competing distractor items are considered and discarded. This is necessary in the Search condition of the present experiment, and possibly also in the Pop-out condition albeit to a lesser extent. It therefore comes as no surprise that we observed robust activation across the lateral frontal lobe (Figures 2 and 4).

Importantly, we observed differences in the proportional activation across subregions of lateral frontal cortex, whereby the IFG was most strongly active, followed by MFG and SFG, in a gradient of increasing activation from ventral to dorsal subregions. These observations may be interpreted from the perspective of preferential engagement of the ventral attention network in this experiment (Corbetta & Shulman, 2002; Corbetta et al., 2008). From the perspective of working memory and PFC, several hypotheses exist for the functional specialization of PFC subregions (reviewed in Curtis & D’Esposito, 2003). Some theories make a “material-specific” distinction whereby dorsal subregions are involved in the maintenance of *spatial* information, while ventral subregions are responsible for the maintenance of *object-related* information. Other models make a “process-specific” distinction whereby dorsal subregions support higher-order control functions like manipulation of items in working memory, while ventral subregions are engaged in “simpler” operations, such as encoding and retrieval of items into and from memory. Our data cannot disentangle these propositions. Our data does not support a strict dichotomy between dorsal and ventral subregions of lateral prefrontal cortex, as we observe a continuum of increasing activity from the superior to middle to inferior frontal gyri in this classical visual search task.

We observed proportionally less activation in the OFC region (Figure 4). This is consistent with a vast set of work documenting different roles for the lateral prefrontal cortex versus the OFC in behavior (reviewed in Hartikainen & Knight, 2003). Ventral frontal regions have a prominent role in the default mode network (Shulman et al., 1997; Raichle et al., 2001; Raichle, 2015). Hence, we might expect to find decreases rather than increases in the OFC in this task (Ossandon et al., 2011; Raccah et al., 2018). Lateral temporal cortex (STG and STS), also a component of the default mode network, similarly showed a very low proportional activation rate (Figure 5).

#### The medial temporal lobe is engaged on par with frontal and parietal cortex

We observed robust MTL activation during the search interval (Figures 2 and 3) in this experiment: The proportion active electrodes in the MTL was indistinguishable from the frontal and parietal cortices. We detected active electrode sites in entorhinal cortex, hippocampus and, to a lesser extent, the parahippocampal cortex. This result is consistent with previous work demonstrating a role for the entorhinal grid cell system in free-viewing of natural scenes in stationary macaques (Killian et al., 2012; Killian et al., 2015). This raises the possibility that the grid cell system might also be engaged in this simpler and more artificial task scene analysis. The result is also consistent with research demonstrating visual working memory (Olson et al., 2006) and implicit visuospatial memory (Chun & Phelps, 1999) impairments in patients with MTL lesions. To the best of our knowledge, this is the first time that MTL engagement, including hippocampus and entorhinal cortex, has been demonstrated in stationary humans performing a controlled, classical visual search paradigm. One previous study has reported individual target-selective cells in the human hippocampus and amygdala while searching a display of naturalistic images (Wang et al., 2018). No equivalent result has been reported, however, which includes the entorhinal cortex in a classical Search versus Pop-out paradigm. The implication is that visual attention work should add MTL structures to the well-known cortical ROIs, such as the IPs, FEF, and dorsolateral PFC (dlPFC), when studying top-down and bottom-up visual attention in search. Classical “attention” effects in serial search and pop-out may need to be re-examined through the lens of viewing these search behaviors as navigation behaviors in visual space (Meister & Buffalo, 2016, Nau et al., 2018).

Adjacent to the MTL, we observed an activation rate of 37% in the amygdala. This effect may be driven by its role in detecting novelty and salience (Rutishauser et al., 2006); the observation that its activity is modulated by visual fixations in free-viewing natural images (Minxha et al., 2017); and its role in a broader limbic network including the MTL.

In sum, this work confirms and extends a central role for the frontoparietal attention network in visual search. We observed prominent regional differences within the PFC, highlighting clear distinctions between OFC versus lateral PFC, and a graded activation profile across lateral PFC. Moreover, we provide support for a central role for the MTL, including the entorhinal cortex and the amygdala, in this classical Search versus Pop-out paradigm. Visual search in humans appears to engage widely distributed brain regions and is not restricted to isolated regions in the parietal and frontal cortices as previously reported (Buschman & Miller, 2007; Li et al., 2010).

### Condition-selective effects

#### Most task-active sites are equally engaged in Pop-out and Search

We find that the vast majority of task-active sites do not show significant condition modulations (91.7%), consistent with past research documenting a largely overlapping network of regions in the frontal and parietal cortices, engaged in both tasks (Leonards et al., 2000; Ossandon et al., 2012; Katsuki & Constantinidis, 2014). The observation of substantial shared neural infrastructure in these two visual search tasks may be influenced by low-level feature detection (edges and colors), object detection, decision-making (whether to select the left or right response), motor planning, etc. We nonetheless detected a sparse, yet robust, set of condition-selective sites. We describe their anatomical distributions below.

#### Search-selective sites are concentrated in lateral frontal cortex

Consistent with a framework that proposes that frontal cortex has a critical role in facilitating Search as opposed to Pop-out, (Li et al., 2010; Buschman & Miller, 2007), we find Search-selective sites exclusively in the frontal and cingulate cortices. Specifically, the majority of Search-selective electrodes were located in lateral frontal cortex. While the number of electrodes (n=9) showing significant Search-selective effects were few, they occurred across multiple patients with PFC coverage (N=4), and the effects were among the largest we observed in this dataset (see Figure 6.C).

This result is consistent with numerous past studies, reporting increased lateral frontal cortex engagement in Search but not Pop-out across a range of recording modalities. fMRI studies have shown increased BOLD signal in regions of lateral frontal cortex in Search tasks relative to Pop-out (SFS: Leonards et al., 2000; MFG and IFG: Anderson et al., 2007; 2010). Other fMRI work has shown a more general role for lateral frontal cortex (MFG) in guiding top-down visual attention (Gazzaley et al., 2007).

In a macaque lesion study, Rossi et al. (2007) showed that ablating the lateral surface of right PFC impaired search performance in a difficult search condition, in which the search cue was often switched across trials, but not in a color pop-out condition similar to the present experiment. Partially convergent evidence was reported by Iba and Sawaguchi (2003), who found that a muscimol injection to dlPFC of rhesus macaques caused an equal impairment to a conjunction (difficult) and pop-out search.

TMS studies in humans have, similarly, demonstrated a causal role for dlPFC in enabling performance in Search, but not Pop-out (Kalla et al., 2009). One of these TMS studies was conducted on an identical Search and Pop-out paradigm as the one employed in the present study (Yan et al., 2016).

A plausible functional explanation for this preferential engagement of lateral prefrontal cortex in Search is its greater demands on working memory as compared to Pop-out. Indeed, Anderson and colleagues (2010) used a joint working memory (verbal and spatial) and visual search paradigm to demonstrate that the right MFG and IFG were engaged both in working memory and Search. They also showed that, on a behavioral level, simultaneous engagement in a working memory task, impaired serial search performance. Hence, our results, showing sparse but robust preferential engagement of the lateral frontal cortex in Search, fit into the extant literature. This body of research converges to demonstrate that Search places greater demands on working memory, and these greater working memory demands are reflected in neural activity in lateral frontal cortex.

#### Pop-out selective sites are distributed across the cortical hierarchy, and include lateral frontal cortex

A number of brain regions have been proposed to be the most critical area for visual pop-out. Several empirical studies highlight a central role for parietal cortex, especially areas adjacent to the IPs (Gottlieb et al., 1998; Buschman & Miller, 2007; Li et al., 2010; Wardak et al., 2010; Yan et al., 2016).

In contrast, a prominent computational theory (Zhaoping, 2002; Zhaoping & Dayan, 2006; Zhaoping, 2019) and related neural evidence from macaques (Yan et al., 2018), emphasize the importance of early salience computations in primary visual cortex (V1). Zhaoping’s theory incorporates the possibility of feedback from higher-order visual areas, such as V2, V3, V4, and IT cortex (Zhaoping, 2019). It does not, however, include any putative involvement of higher-order cortical areas, such as the parietal or frontal cortices. It should be noted that salience maps have been observed in several cortical areas beyond V1, including: area V4 in the occipital lobe (Burrows & Moore, 2009), area LIP in the parietal cortex (Gottlieb et al., 1998), and the FEF in the frontal lobe (Thompson & Bichot, 2005). Visual salience effects have also been reported in subcortical regions including the pulvinar nucleus of the thalamus (Robinson & Petersen, 1992), and the superior colliculus (White et al., 2017).

In sharp contrast to these bottom-up theories, which depict visual pop-out as reflecting anatomically early detection of visual salience, Hochstein and Ahissar (2002), proposed the Reverse Hierarchy Theory (RHT) for vision. According to this theory, the visual pop-out phenomenon should originate in high-order cortical areas such as frontal cortex, where large receptive fields can account for the broad spread of visual attention necessary for parallel detection of a salient object. This is consistent with viewing Pop-out as a high-level object detection task (Hochstein & Ahissar 2002; see also Nakayama & Martini, 2011), wherein the pop-out effect is invariant to object size on the retina, unlike the view of Pop-out as detection of low-level feature anomalies based on lateral inhibition in V1 (Zhaoping, 2002; Zhaoping & Dayan, 2006; Zhaoping, 2019). Consistent with RHT, two macaque studies using fMRI and single-neuron recordings respectively, reported lateral frontal cortex responses to visual pop-out (Wardak et al., 2010, using fMRI; Katsuki & Constantinidis, 2012, using single-neuron recordings).

Finally, a theory for cortical engagement in visual pop-out has been proposed by Corbetta & Shulman (2002), and extended by Corbetta et al. (2008). This theory proposes that bottom-up capture of attention by a salient stimulus, as in visual pop-out, is accomplished by a distributed network of cortical regions, notably all in the right hemisphere, including the TPJ as well as parts of the MFG, IFG, frontal operculum and anterior insula (the right-lateralized ventral attention network, VAN).

Our data reveal a distributed set of sites selective for Pop-out, in partial agreement with each of the extant theories. We observed several Pop-out selective sites in the parietal lobe as predicted by past research though not all were adjacent to the IPs. A single electrode site in the occipital lobe was selective for Pop-out. We refrain from interpreting this result in light of the limited coverage in this area. Temporal lobe Pop-out selectivity can be understood from the perspective of the importance of ventral temporal regions for object detection (DiCarlo & Cox, 2007; DiCarlo et al., 2012). Similarly, Pop-out selectivity in ventral temporal cortex has previously been observed in other color pop-out paradigms (Wardak et al., 2010; Ossandon et al., 2012).

Our most striking finding is the presence of Pop-out selective sites in frontal cortex. These sites were detected not only in areas at or near the FEF, which has a known role in detecting salience and planning eye movements, but also in lateral frontal cortex and OFC. This result is consistent with the view that top-down processing has a prominent role in implementing visual pop-out (Hochstein & Ahissar, 2002; Nakayama & Martini, 2011). It is inconsistent with a view of visual pop-out as exclusively implemented in V1 and adjacent occipital lobe areas (Zhaoping, 2002; Zhaoping & Dayan, 2006; Zhaoping, 2019). Our result may explain Iba and Sawaguchi’s (2003) observation that a muscimol injection to dlPFC of monkeys caused impairments in both pop-out and serial search. Future causal investigations of the role of lateral frontal cortex in Pop-out and Search may find differential impairment in the two conditions if the sites of the temporary lesion or stimulation is selected with greater anatomical precision within dlPFC than has been possible in the past. Furthermore, our result is consistent with two previous functional studies in macaques which also reported the presence of Pop-out selective sites in lateral frontal cortex (Wardak et al., 2010; Katsuki & Constantinidis, 2012). To the best of our knowledge, this is the first time that such sites have been detected in humans.

While our data is consistent with the view that Pop-out is implemented in a distributed set of sites across cortex, they do not fit neatly with the framework positing a right-lateralized ventral attention network for salience-driven orienting of attention (Corbetta & Shulman, 2002; Corbetta et al., 2008), since we also observe robust Pop-out selective effects in the left hemisphere (Figure 6). In sum, our results do not support exclusivist claims about regional engagement in Pop-out, such as the view that Pop-out is solely a bottom-up phenomenon, not engaging frontal cortex. We have demonstrated that the known presence of salience maps in low-level visual regions, as well as parietal cortex and FEF, is not mutually exclusive with the simultaneous rapid engagement of a high-level cognitive region, the lateral frontal cortex, in visual pop-out.

## Conclusion

The present study documents two previously unreported neural mechanisms subserving visual search and pop-out in humans. First lateral PFC engagement shows regional differences with greater activation in inferior than superior regions, and diminished proportional activations in OFC. Second, we establish a central role for the medial temporal lobe, including the hippocampus, entorhinal cortex, and amygdala, in visual search and pop-out in humans. We additionally confirm a central role for lateral PFC in serial search. We observe a distributed set of Pop-out selective sites across cortex. In sum, this study provides evidence for a more distributed processing view of visual attention.

## Acknowledgments

We thank each of the patients, who volunteered their time to take part in this experiment. We thank Ling Li for sharing an earlier version of the experiment (Li et al., 2010; 2013), James Lubell for assistance with experiment programming, and Jerren Chiu for assistance with data preprocessing. We thank Michael Silver, Athina Tzovara, Lucy Stephenson, and Michael Eickenberg for helpful comments on earlier versions of this manuscript. We thank Anaïs Llorens, Julia Kam, and Avgusta Shestyuk for helpful discussion. We thank the recording team at each hospital for their help with data collection. This work was supported by NINDS R3721135 and NIMH CONTE Center 1P50 MH109529-01 (to R.T.K.), the Research Council of Norway 240389/F20 and Internal Funding from the University of Oslo (to A.-K.S., T.E. and P.G.L.), and UC Irvine Bridge Fund (to J.J.L.)

## Author contributions

S.J.K.S. adapted the experiment for intracranial recording, supervised the data collection, analysed the data, and wrote the paper. R.T.K. conceived of the project and supervised its development. S.S. and S.J.K.S. devised the statistical testing approach for task-selective and condition-selective electrodes. R.J. created the brain reconstructions in Figures 2 and 6. D.K.-S., K.D.L., P.B.W., T.E., P.G.L., A.-K.S., and J.J.L. contributed to data accessibility and revision of the manuscript.

## References

Anderson, E. J., Mannan, S. K., Husain, M., Rees, G., Sumner, P., Mort, D. J., McRobbie, D., Kennard, C. (2007). Involvement of prefrontal cortex in visual search. Experimental Brain Research, 180(2), 289–302. https://doi.org/10.1007/s00221-007-0860-0

Anderson, E. J., Mannan, S. K., Rees, G., Sumner, P., & Kennard, C. (2010). Overlapping functional anatomy for working memory and visual search. Experimental Brain Research, 200(1), 91–107. https://doi.org/10.1007/s00221-009-2000-5

Barceló, F., Suwazono, S., & Knight, R. T. (2000). Prefrontal modulation of visual processing in humans. Nature Neuroscience, 3(4), 399–403.

Burrows, B. E., & Moore, T. (2009). Influence and limitations of popout in the selection of salient visual stimuli by area V4 neurons. Journal of Neuroscience, 29(48), 15169–15177. https://doi.org/10.1523/JNEUROSCI.3710-09.2009

Buschman, T. J., & Miller, E. K. (2007). Top-down versus bottom-up control of attention in the prefrontal and posterior parietal cortices. Science, 315, 1860–1862. https://doi.org/10.1126/science.1138071

Burke, J. F., Zaghloul, K. A., Jacobs, J., Williams, R. B., Sperling, M. R., Sharan, A. D., & Kahana, M. J. (2013). Synchronous and asynchronous theta and gamma activity during episodic memory formation. Journal of Neuroscience, 33(1), 292–304. https://doi.org/10.1523/jneurosci.2057-12.2013

Cauchoix, M., Crouzet, S. M., Fize, D., & Serre, T. (2016). Fast ventral stream neural activity enables rapid visual categorization. NeuroImage, 125(33), 280–290. https://doi.org/10.1016/j.neuroimage.2015.10.012

Chun, M. M., & Phelps, E. A. (1999). Memory deficits for implicit contextual information in human amnesics. Nature Neuroscience, 2(9), 844–847. https://doi.org/10.1038/12222

Collins, D. L., Zijdenbos, A. P., Kollokian, V., Sled, J. G., Kabani, N. J., Holmes, C. J., & Evans, A. C. (1998). Design and Construction of a Realistic Digital Brain Phantom. IEEE Transactions on Medical Imaging, 17(3), 463–468.

Corbetta, M., Patel, G., & Shulman, G. L. (2008). The reorienting system of the human brain: from environment to theory of mind. Neuron, 58(3), 306–324. https://doi.org/10.1016/j.neuron.2008.04.017

Corbetta, M., & Shulman, G. L. (2002). Control of goal-directed and stimulus-driven attention in the brain. Nature Reviews Neuroscience, 3, 201–215. https://doi.org/10.1038/nrn755

Curtis, C. E., & D’Esposito, M. (2003). Persistent activity in the prefrontal cortex during working memory. Trends in Cognitive Sciences, 7(9), 415–423. https://doi.org/10.1016/S1364-6613(03)00197-9

Dale, A. M., Fischl, B., & Sereno, M. I. (1999). Cortical Surface-Based Analysis: I. Segmentation and Surface Reconstruction. NeuroImage, 9(2), 179–194. https://doi.org/https://doi.org/10.1006/nimg.1998.0395

DiCarlo, J. J., & Cox, D. D. (2007). Untangling invariant object recognition. Trends in Cognitive Sciences, 11(8), 333–341. https://doi.org/10.1016/j.tics.2007.06.010

DiCarlo, J. J., Zoccolan, D., & Rust, N. C. (2012). How does the brain solve visual object recognition? Neuron, 73(3), 415–434. https://doi.org/10.1016/j.neuron.2012.01.010

Dykstra, A. R., Chan, A. M., Quinn, B. T., Zepeda, R., Keller, C. J., Cormier, J., Madsen, J. R., Eskandar, E. N., Cash, S. S. (2012). Individualized localization and cortical surface-based registration of intracranial electrodes. NeuroImage, 59, 3563–3570. https://doi.org/10.1016/j.neuroimage.2011.11.046

Eimer, M. (2014). The neural basis of attentional control in visual search. Trends in Cognitive Sciences, 18(10), 526–535. https://doi.org/https://doi.org/10.1016/j.tics.2014.05.005

Gottlieb, J. P., Kusunoki, M., & Goldberg, M. E. (1998). The representation of visual salience in monkey parietal cortex. Nature, 391(6666), 481–484. https://doi.org/10.1038/35135

Gazzaley, A., Rissman, J., Cooney, J., Rutman, A., Seibert, T., Clapp, W., & D’Esposito, M. (2007). Functional interactions between prefrontal and visual association cortex contribute to top-down modulation of visual processing. Cerebral Cortex, 17(SUPPL. 1), 125–135. https://doi.org/10.1093/cercor/bhm113

Haller, M., Case, J., Crone, N. E., Chang, E. F., King-Stephens, D., Laxer, K. D., Weber, P. B., Parvizi, J., Knight, R. T., & Shestyuk, A. Y. (2018). Persistent neuronal activity in human prefrontal cortex links perception and action. Nature Human Behavior, 2(1), 80–91. https://doi.org/10.1038/s41562-017-0267-2.Persistent

Hartikainen, K. M., & Knight, R. T. (2003). Lateral and Orbital Prefrontal Cortex Contributions to Attention. In J. Polich (Ed.), Detection of Change (pp. 99–116). New York: Springer Science and Business Media. https://doi.org/10.1007/978-1-4615-0294-4_6

Hermes, D., Miller, K. J., Noordmans, H. J., Vansteensel, M. J., & Ramsey, N. F. (2010). Automated electrocorticographic electrode localization on individually rendered brain surfaces. Journal of Neuroscience Methods, 185, 293–298. https://doi.org/10.1016/j.jneumeth.2009.10.005

Hochstein, S., & Ahissar, M. (2002). View from the Top: Hierarchies and Reverse Hierarchies in the Visual System. Neuron, 36, 791–804.

Iba, M., & Sawaguchi, T. (2003). Involvement of the dorsolateral prefrontal cortex of monkeys in visuospatial target selection. Journal of Neurophysiology, 89(1), 587–599. https://doi.org/10.1152/jn.00148.2002

Jafarpour, A., Griffin, S., Lin, J. J., & Knight, R. T. (2019). Medial Orbitofrontal Cortex, Dorsolateral Prefrontal Cortex, and Hippocampus Differentially Represent the Event Saliency. Journal of Cognitive Neuroscience, 31(6), 874–884. https://doi.org/10.1162/jocn

Julesz, B. (1981). Textons, the elements of texture perception, and their interactions. Nature, 290(5802), 91–97. https://doi.org/10.1038/290091a0

Kalla, R., Muggleton, N. G., Cowey, A., & Walsh, V. (2009). Human dorsolateral prefrontal cortex is involved in visual search for conjunctions but not features: A theta TMS study. Cortex, 45(9), 1085–1090. https://doi.org/10.1016/j.cortex.2009.01.005

Kastner, S., DeSimone, K., Konen, C. S., Szczepanski, S. M., Weiner, K. S., & Schneider, K. A. (2007). Topographic maps in human frontal cortex revealed in memory-guided saccade and spatial working-memory tasks. Journal of Neurophysiology, 97(5), 3494–3507. https://doi.org/10.1152/jn.00010.2007

Katsuki, F., & Constantinidis, C. (2012). Early involvement of prefrontal cortex in visual bottom up attention. Nature Neuroscience, 15(8), 1160–1168. http://doi.org/10.1038/nn.3164

Katsuki, F., & Constantinidis, C. (2014). Bottom-up and top-down attention: Different processes and overlapping neural systems. Neuroscientist, 20(5), 509–521. https://doi.org/10.1177/1073858413514136

Killian, N. J., Jutras, M. J., & Buffalo, E. A. (2012). A map of visual space in the primate entorhinal cortex. Nature, 491(7426), 761–764. https://doi.org/10.1038/nature11587

Killian, N. J., Potter, S. M., & Buffalo, E. A. (2015). Saccade direction encoding in the primate entorhinal cortex during visual exploration. Proceedings of the National Academy of Sciences, 201417059. https://doi.org/10.1073/pnas.1417059112

Knight, R. T. (1984). Decreased response to novel stimuli after prefrontal lesions in man. Electroencephalography and Clinical Neurophysiology, 59, 9–20.

Knight, R. T., Grabowecky, M. F., & Scabini, D. (1995). Role of human prefrontal cortex in attention control. In H. H. Jasper, S. Riggio, & P. S. Goldman-Rakic (Eds.), Epilepsy and the functional anatomy of the frontal lobe (pp. 21–31). New York: Raven Press.

Knight, R. T. (1996). Contribution of human hippocampal region to novelty detection. Nature, 383, 256–259.

Leonards, U., Sunaert, S., Van Hecke, P., & Orban, G. A. (2000). Attention mechanisms in visual search - An fMRI study. Journal of Cognitive Neuroscience, 12 (SUPPL. 2), 61–75. http://doi.org/10.1162/089892900564073

Li, L., Gratton, C., Fabiani, M., & Knight, R. T. (2013). Age-related frontoparietal changes during the control of bottom-up and top-down attention: An ERP study. Neurobiology of Aging, 34, 477–488. https://doi.org/10.1016/j.neurobiolaging.2012.02.025

Li, L., Gratton, C., Yao, D., & Knight, R. T. (2010). Role of frontal and parietal cortices in the control of bottom-up and top-down attention in humans. Brain Research, 1344, 173–184. https://doi.org/10.1016/j.brainres.2010.05.016

Logothetis, N. K., Pauls, J., Augath, M., Trinath, T., & Oeltermann, A. (2001). Neurophysiological investigation of the basis of the fMRI signal. Nature, 412, 150–157. https://doi.org/10.1038/35084005

Mackey, W. E., Winawer, J., & Curtis, C. E. (2017). Visual field map clusters in human frontoparietal cortex. ELife, 6, 1–23. https://doi.org/10.7554/elife.22974

Mangano, G. R., Oliveri, M., Turriziani, P., Smirni, D., Zhaoping, L., & Cipolotti, L. (2015). Repetitive transcranial magnetic stimulation over the left parietal cortex facilitates visual search for a letter among its mirror images. Neuropsychologia, 70, 196–205. https://doi.org/10.1016/j.neuropsychologia.2015.03.002

Maris, E., & Oostenveld, R. (2007). Nonparametric statistical testing of EEG- and MEG-data. Journal of Neuroscience Methods, 164(1), 177–190. https://doi.org/10.1016/j.jneumeth.2007.03.024

Martin, A. B., Yang, X., Saalmann, Y. B., Wang, L., Shestyuk, A., Lin, J. J., Parvizi, J., Knight, R. T., & Kastner, S. (2019). Temporal Dynamics and Response Modulation across the Human Visual System in a Spatial Attention Task : An ECoG Study, 39(2), 333–352.

Meister, M. L. R., & Buffalo, E. A. (2016). Getting directions from the hippocampus: The neural connection between looking and memory. Neurobiology of Learning and Memory, 134, 135–144. https://doi.org/https://doi.org/10.1016/j.nlm.2015.12.004

Minxha, J., Mosher, C., Morrow, J. K., Mamelak, A. N., Adolphs, R., Gothard, K. M., & Rutishauser, U. (2017). Fixations Gate Species-Specific Responses to Free Viewing of Faces in the Human and Macaque Amygdala. Cell Reports, 18(4), 878–891. https://doi.org/10.1016/j.celrep.2016.12.083

Mukamel, R., Gelbard, H., Arieli, A., Hasson, U., Fried, I., & Malach, R. (2005). Coupling between neuronal firing, field potentials, and fMRI in human auditory cortex. Science, 309, 951–954. https://doi.org/10.1126/science.1110913

Nakayama, K., & Martini, P. (2011). Situating visual search. Vision Research, 51(13), 1526–1537. https://doi.org/10.1016/j.visres.2010.09.003

Nir, Y., Fisch, L., Mukamel, R., Gelbard-Sagiv, H., Arieli, A., Fried, I., & Malach, R. (2007). Coupling between Neuronal Firing Rate, Gamma LFP, and BOLD fMRI Is Related to Interneuronal Correlations. Current Biology, 17, 1275–1285. https://doi.org/10.1016/j.cub.2007.06.066

Nobre, A. C., Coull, J. T., Walsh, V., & Frith, C. D. (2002). Brain Activations during Visual Search: Contributions of Search Efficiency versus Feature Binding. NeuroImage, 18(1), 91–103. https://doi.org/https://doi.org/10.1006/nimg.2002.1329

Olson, I. R., Moore, K. S., Stark, M., & Chatterjee, A. (2006). Visual working memory is impaired when the medial temporal lobe is damaged. Journal of Cognitive Neuroscience, 18(7), 1087–1097. https://doi.org/10.1162/jocn.2006.18.7.1087

Oostenveld, R., Fries, P., Maris, E., & Schoffelen, J. M. (2011). FieldTrip: Open source software for advanced analysis of MEG, EEG, and invasive electrophysiological data. Computational Intelligence and Neuroscience, 2011. https://doi.org/10.1155/2011/156869

Ossandon, T., Jerbi, K., Vidal, J. R., Bayle, D. J., Henaff, M.-A., Jung, J., Minotti, L., Bertrand, O., Kahane, P., & Lachaux, J.-P. (2011). Transient Suppression of Broadband Gamma Power in the Default-Mode Network Is Correlated with Task Complexity and Subject Performance. The Journal of Neuroscience, 31(41), 14521–14530. https://doi.org/10.1523/JNEUROSCI.2483-11.2011

Ossandon, T., Vidal, J. R., Ciumas, C., Jerbi, K., Hamame, C. M., Dalal, S. S., Bertrand, O., Minotti, L., Kahane, P., & Lachaux, J.-P. (2012). Efficient “Pop-Out” Visual Search Elicits Sustained Broadband Gamma Activity in the Dorsal Attention Network. Journal of Neuroscience, 32(10), 3414–3421. https://doi.org/10.1523/JNEUROSCI.6048-11.2012

Parker, A., Wilding, E., & Akerman, C. (1998). The von Restorff effect in visual object recognition memory in humans and monkeys: The role of frontal/perirhinal interaction. Journal of Cognitive Neuroscience, 10(6), 691–703. https://doi.org/10.1162/089892998563103

Parvizi, J., & Kastner, S. (2018). Promises and limitations of human intracranial electroencephalography. Nature Neuroscience, 21(4), 474–483. https://doi.org/10.1038/s41593-018-0108-2

Raichle, M. E., MacLeod, A. M., Snyder, A. Z., Powers, W. J., Gusnard, D. A., & Shulman, G. L. (2001). A default mode of brain function. PNAS, 98(2), 676–682.

Raichle, M. E. (2015). The Brain’s Default Mode Network. Annual Review of Neuroscience, 38(1), 433–447. https://doi.org/10.1146/annurev-neuro-071013-014030

Ray, S., Crone, N. E., Niebur, E., Franaszczuk, P. J., & Hsiao, S. S. (2008). Neural correlates of high-gamma oscillations (60-200 Hz) in macaque local field potentials and their potential implications in electrocorticography. The Journal of Neuroscience, 28(45), 11526–11536. https://doi.org/10.1523/JNEUROSCI.2848-08.2008

Ray, S., & Maunsell, J. H. R. (2011). Different origins of gamma rhythm and high-gamma activity in macaque visual cortex. PLoS Biology, 9(4), 1–15. https://doi.org/10.1371/journal.pbio.1000610

Regev, T. I., Winawer, J., Gerber, E. M., Knight, R. T., & Deouell, L. Y. (2018). Human posterior parietal cortex responds to visual stimuli as early as peristriate occipital cortex. European Journal of Neuroscience (Vol. 48). https://doi.org/10.1111/ejn.14164

von Restorff, H. (1933) Psychol. Forsch. 18: 299. https://doi.org/10.1007/BF02409636

Robinson, D. L., & Petersen, S. E. (1992). The pulvinar and visual salience. Trends in Neurosciences, 15(4), 127–132.

Rossi, A. F., Bichot, N. P., Desimone, R., & Ungerleider, L. G. (2007). Top Down Attentional Deficits in Macaques with Lesions of Lateral Prefrontal Cortex. Journal of Neuroscience, 27(42), 11306–11314. https://doi.org/10.1523/JNEUROSCI.2939-07.2007

Rutishauser, U., Mamelak, A. N., & Schuman, E. M. (2006). Single-trial learning of novel stimuli by individual neurons of the human hippocampus-amygdala complex. Neuron, 49(6), 805–813. https://doi.org/10.1016/j.neuron.2006.02.015

Shulman, G. L., Fiez, J. A., Corbetta, M., Buckner, R. L., Miezin, F. M., Raichle, M. E., & Petersen, S. E. (1997). Common blood flow changes across visual tasks: II. Decreases in cerebral cortex. Journal of Cognitive Neuroscience, 9(5), 648–663. https://doi.org/10.1162/jocn.1997.9.5.648

Silver, M. A., Ress, D., & Heeger, D. J. (2005). Topographic maps of visual spatial attention in human parietal cortex. Journal of Neurophysiology, 94(2), 1358–1371. https://doi.org/10.1152/jn.01316.2004

Silver, M. A., & Kastner, S. (2009). Topographic maps in human frontal and parietal cortex. Trends in Cognitive Sciences, 13(11), 488–495. https://doi.org/10.1016/j.tics.2009.08.005.

Stolk, A., Griffin, S., Van Der Meij, R., Dewar, C., Saez, I., Lin, J. J., Piantoni, G., Schoffelen, J. M., Knight, R. T., Oostenveld, R. (2018). Integrated analysis of anatomical and electrophysiological human intracranial data. Nature Protocols, 13(7), 1699–1723. https://doi.org/10.1038/s41596-018-0009-6

Thompson, K. G., & Bichot, N. P. (2005). A visual salience map in the primate frontal eye field. Progress in Brain Research. https://doi.org/10.1016/S0079-6123(04)47019-8

Treisman, A. M., & Gelade, G. (1980). A feature-integration theory of attention. Cognitive Psychology, 12, 97–136. Retrieved from http://www.ncbi.nlm.nih.gov/pubmed/7351125

Wang, S., Mamelak, A. N., Adolphs, R., & Rutishauser, U. (2018). Encoding of Target Detection during Visual Search by Single Neurons in the Human Brain. Current Biology, 28(13), 2058–2069. https://doi.org/https://doi.org/10.1016/j.cub.2018.04.092

Wardak, C., Vanduffel, W., & Orban, G. A. (2010). Searching for a salient target involves frontal regions. Cerebral Cortex, 20(10), 2464–2477. https://doi.org/10.1093/cercor/bhp315

White, B. J., Kan, J. Y., Levy, R., Itti, L., & Munoz, D. P. (2017). Superior colliculus encodes visual saliency before the primary visual cortex. Proceedings of the National Academy of Sciences of the United States of America, 114(35), 9451–9456. https://doi.org/10.1073/pnas.1701003114

Wolfe, J. (2014). Approaches to Visual Search: Feature Integration Theory and Guided Search. In S. Kastner & A. C. Nobre (Eds.), The Oxford Handbook of Attention (1st ed., pp. 11–55). Oxford, United Kingdom: Oxford University Press.

Wolfe, J. M. (2018). Visual Search. In J. T. Wixted (Ed.), Stevens’ Handbook of Experimental Psychology and Cognitive Neuroscience (4th ed., pp. 1–55). John Wiley & Sons, Inc. http://doi.org/10.1002/9781119170174.epcn213

Xu, Y. (2018). The Posterior Parietal Cortex in Adaptive Visual Processing. Trends in Neurosciences, 41(11), 806–822. https://doi.org/10.1016/j.tins.2018.07.012

Yan, Y., Wei, R., Zhang, Q., Jin, Z., & Li, L. (2016). Differential roles of the dorsal prefrontal and posterior parietal cortices in visual search: A TMS study. Scientific Reports, 6(January), 1–9. https://doi.org/10.1038/srep30300

Yan, Y., Zhaoping, L., & Lia, W. (2018). Bottom-up saliency and top-down learning in the primary visual cortex of monkeys. Proceedings of the National Academy of Sciences of the United States of America, 115(41), 10499–10504. https://doi.org/10.1073/pnas.1803854115

Zhaoping, L. (2002). A saliency map in primary visual cortex. Trends in Cognitive Sciences, 6(1), 9–16.

Zhaoping, L., & Dayan, P. (2006). Pre-attentive visual selection. Neural Networks, 19(9), 1437–1439. https://doi.org/10.1016/j.neunet.2006.09.003

Zhaoping, L. (2019). A new framework for understanding vision from the perspective of the primary visual cortex. Current Opinion in Neurobiology, 58(Box 1), 1–10. https://doi.org/10.1016/j.conb.2019.06.001

