## Supplementary Tables and Figures for "Intracranial recordings demonstrate medial temporal lobe engagement in visual search in humans"

**Supplementary Information**

**Supplementary Table 1.** Participant information.

| SID | Sex | Age | Ha. | SEEG/ ECoG | He. | ROIs | Number Included contacts | Number active contacts | Number included (compl.) trials Pop-Out | Number included. (compl.) trials Search |
|---|---|---|---|---|---|---|---|---|---|---|
| S1 | F | 28 | R | ECoG | R+L | T, MTL | 22 | 5 | 57 (64) | 53 (64) |
| S2 | F | 63 | R | SEEG | R+L | T, F, MTL, cing., amy. | 51 | 19 | 48 (64) | 43 (64) |
| S3 | M | 27 | R | SEEG | R+L | F, P, T, MTL, amy. | 40 | 7 | 51 (64) | 50 (64) |
| S4 | F | 34 | R | SEEG | R+L | F, T, MTL, cing., amy. | 61 | 18 | 48 (67) | 38 (61) |
| S5 | M | 35 | R | SEEG | R | F, cing. | 36 | 17 | 58 (64) | 42 (64) |
| S6 | M | 52 | R | SEEG | R | F, P, cing. | 75 | 33 | 40 (50) | 38 (46) |
| S7 | F | 23 | R | ECoG | R+L | F, T | 54 | 17 | 59 (64) | 56 (64) |
| S8 | F | 36 | R | ECoG | R+L | F, P, cing. | 48 | 20 | 56 (64) | 52 (64) |
| S9 | M | 32 | R | ECoG | R+L | F, T, MTL, cing. | 53 | 12 | 49 (64) | 33 (64) |
| S10 | F | 69 | R | ECoG | R+L | F, P, T, MTL | 35 | 11 | 59 (64) | 29 (64) |
| S11 | M | 25 | A | ECoG | R+L | F, P, T, MTL, cing. | 59 | 11 | 53 (64) | 42 (64) |
| S12 | F | 21 | R | ECoG | R+L | F, P, cing. | 107 | 47 | 57 (64) | 49 (64) |
| S13 | F | 58 | R | SEEG | R+L | F, T, MTL, cing., amy. | 53 | 35 | 57 (64) | 50 (64) |
| S14 | M | 34 | A | SEEG | R+L | F, T, MTL, cing., amy. | 37 | 5 | 62 (64) | 60 (64) |
| S15 | F | 22 | R | SEEG | R+L | F, T, cing. | 65 | 30 | 42 (64) | 30 (64) |
| S16 | M | 41 | R | SEEG | R+L | F, T, MTL, cing., amy. | 49 | 11 | 51 (64) | 38 (64) |

| | | | | | | | | | | |
|---|---|---|---|---|---|---|---|---|---|---|
| S17 | F | 32 | R | SEEG | R+L | F, T, MTL, ACC, amy. | 35 | 8 | 54 (64) | 31 (64) |
| S18 | F | 26 | R | ECoG | L | P, T, O, MTL | 125 | 33 | 58 (64) | 45 (64) |
| S19 | M | 23 | R | SEEG | R+L | F, T, cing. | 78 | 37 | 61 (64) | 60 (64) |
| S20 | F | 50 | R | SEEG | R+L | F, T, MTL, cing. | 31 | 9 | 34 (64) | 21 (64) |
| S21 | M | 27 | R | SEEG | R+L | F, T, cing., amy. | 68 | 15 | 48 (64) | 38 (64) |
| S22 | M | 23 | R | SEEG | R | F, T, O, MTL, cing., amy. | 42 | 13 | 44 (64) | 36 (64) |
| S23 | F | 25 | R | SEEG | R+L | F, T, MTL, cing., amy. | 97 | 42 | 56 (64) | 43 (64) |

Table header: SID, subject ID; Ha., handedness; He., hemispheric coverage; ROIs, coverage by ROI; Number included (compl.) trials, number included (completed) trials.
Sex: F, female; M, male. Handedness: R, right; L, left; A, ambidextrous. Hemispheric coverage: R, right; L, left; R + L, right and left hemispheres. Coverage by ROI: F, frontal; P, parietal; T, temporal; O, occipital; MTL, medial temporal lobe; cing., cingulate; amy., amygdala.

**Supplementary Table 2.** Distribution of task-selective effects, by large cortical ROIs.

| **Anatomical ROI** | **Percent active electrodes (active/included)** | | |
|---|---|---|---|
| | left | right | total |
| Frontal cortex | **44%** (75/171) | **31%** (88/281) | **36%** (163/452) |
| Parietal cortex | **44%** (12/27) | **43%** (23/53) | **44%** (35/80) |
| Temporal cortex | **20%** (46/227) | **23%** (33/142) | **21%** (79/369) |
| Occipital cortex | **73%** (8/11) | **50%** (3/6) | **65%** (11/17) |
| Cingulate cortex | **22%** (16/72) | **28%** (21/74) | **25%** (37/146) |
| Medial temporal lobe | **48%** (15/31) | **33%** (9/27) | **41%** (24/58) |
| Amygdala | **27%** (4/15) | **43%** (10/23) | **37%** (14/38) |
| Sensorimotor cortex | **66%** (57/87) | **45%** (33/74) | **56%** (90/161) |
| Total | **36%** (233/641) | **32%** (220/680) | **34%** (453/1321) |
| **Total excluding sensorimotor cortex** | **32%** (176/554) | **31%** (187/606) | **31%** (363/1160) |

**Supplementary Table 3.** Distribution of task-selective effects within PFC.

| **Frontal ROI** | **Percent active electrodes (active/included)** | | |
|---|---|---|---|
| | left | right | total |
| Superior frontal gyrus and sulcus | **29%** (12/42) | **32%** (28/87) | **31%** (40/129) |
| Middle frontal gyrus | **55%** (32/58) | **33%** (26/78) | **43%** (58/136) |
| Inferior frontal gyrus and sulcus | **68%** (17/25) | **46%** (22/48) | **53%** (39/73) |
| Orbitofrontal cortex | **19%** (6/32) | **19%** (8/43) | **19%** (14/75) |
| Medial prefrontal cortex | **57%** (8/14) | **16%** (4/25) | **31%** (12/39) |
| **Total** | **44%** (75/171) | **31%** (88/281) | **36%** (163/452) |

**Supplementary Table 4.** Distribution of task-selective effects within parietal (non-motor) cortex.

| **Parietal ROI** | **Percent active electrodes (active/included)** | | |
|---|---|---|---|
| | left | right | total |
| Superior parietal lobule | **0%** (0/0) | **65%** (11/17) | **65%** (11/17) |
| Inferior parietal lobule | **38%** (9/24) | **29%** (9/31) | **33%** (18/55) |
| Precuneus | **100%** (3/3) | **60%** (3/5) | **75%** (6/8) |
| **Total** | **44%** (12/27) | **43%** (23/53) | **44%** (35/80) |

**Supplementary Table 5.** Distribution of task-selective effects within temporal cortex.

| **Temporal region of interest** | **Percent active electrodes (active/included)** | | |
|---|---|---|---|
| | left | right | total |
| Insula | **15%** (3/20) | **35%** (9/26) | **26%** (12/46) |
| Superior temporal gyrus | **9%** (3/35) | **11%** (1/9) | **9%** (4/44) |
| Superior temporal sulcus | **9%** (4/44) | **13%** (5/38) | **11%** (9/82) |
| Middle temporal gyrus | **20%** (10/50) | **21%** (6/29) | **20%** (16/79) |
| Inferior temporal gyrus | **23%** (8/35) | **21%** (5/24) | **22%** (13/59) |
| Ventral stream | **42%** (13/31) | **38%** (5/13) | **41%** (18/44) |
| Temporal pole | **42%** (5/12) | **67%** (2/3) | **47%** (7/15) |
| **Total** | **20%** (46/227) | **23%** (33/142) | **21%** (79/369) |

**Supplementary Table 6.** Distribution of task-selective effects within cingulate cortex.

| **Cingulate ROI** | **Percent active electrodes (active/included)** | | |
|---|---|---|---|
| | left | right | total |
| Anterior cingulate cortex | **18%** (6/33) | **12%** (4/34) | **15%** (10/67) |
| Midcingulate cortex | **26%** (7/27) | **44%** (11/25) | **35%** (18/52) |
| Posterior cingulate cortex | **25%** (3/12) | **40%** (6/15) | **33%** (9/27) |
| **Total** | **22%** (16/72) | **28%** (21/74) | **25%** (37/146) |

**Supplementary Table 7.** Distribution of task-selective effects within the medial temporal lobe.

| MTL ROI | Percent active electrodes (active/included) | | |
|---|---|---|---|
| | left | right | total |
| Hippocampus | **38%** (5/13) | **44%** (4/9) | **41%** (9/22) |
| Parahippocampal cortex | **50%** (6/12) | **10%** (1/10) | **32%** (7/22) |
| Entorhinal cortex | **67%** (4/6) | **50%** (4/8) | **57%** (8/14) |
| **Total** | **48%** (15/31) | **33%** (9/27) | **41%** (24/58) |

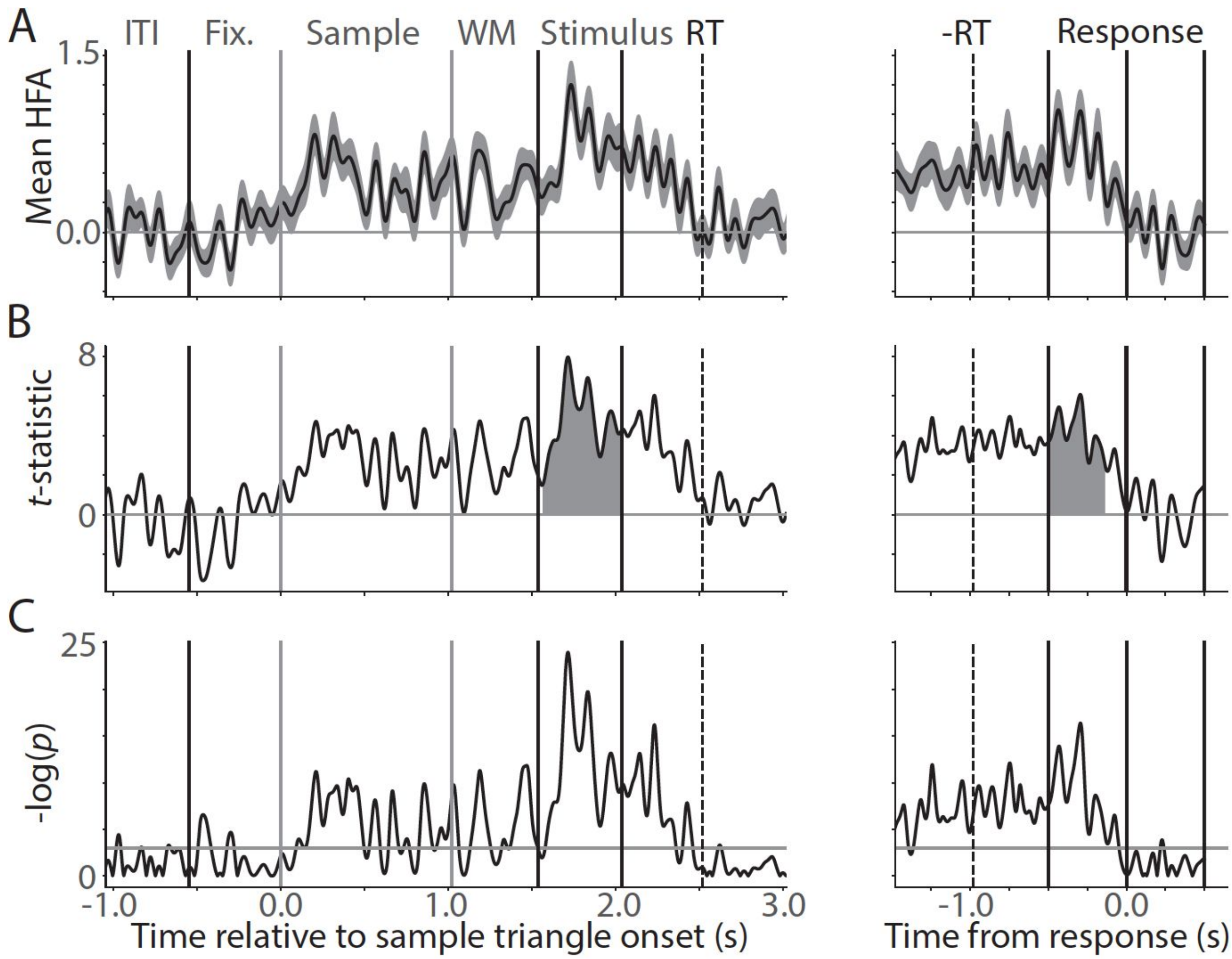


**Supplementary Figure 1.** Example electrode and condition (Search) that was task-selective both in the first considered interval (stimulus onset to 500 ms, Stimulus) and the second considered interval (-500 ms to response). A) Mean, z-scored HFA signal, time-locked to sample triangle onset (left panel), and to the response (right panel). On the left panel, the first vertical line shows the offset of the baseline time-period. The second line shows the onset of the sample triangle. The third line shows the offset of the sample triangle. The fourth line shows stimulus onset. The fifth line shows the offset (500 ms) of the first task interval. The dotted line shows the median RT. On the right panel, the dotted line shows the median stimulus onset time. The second line shows the start of the second task interval (-500 ms relative to response). The third line shows the RT, and the fourth line marks 500 ms after the response. Grey labels indicate intervals between vertical lines. Black labels refer to time points (the vertical lines themselves). ITI, Inter-Trial Interval; Fix., Fixation Interval; Sample, Sample Interval; WM, Working Memory Interval; Stimulus, Stimulus Display Interval; RT, Response Time. B) One-sample *t*-statistics corresponding to the mean HFA traces on top. Shaded areas indicate significant clusters. The area of these clusters were used as the statistics considered in the permutation test. C) -log(*p*) values corresponding to the *t*-statistics in (B). The grey horizontal line shows $p = 0.05$. Any points above this line were significantly greater than the pooled baseline used to z-score the mean signal, as shown in (A).

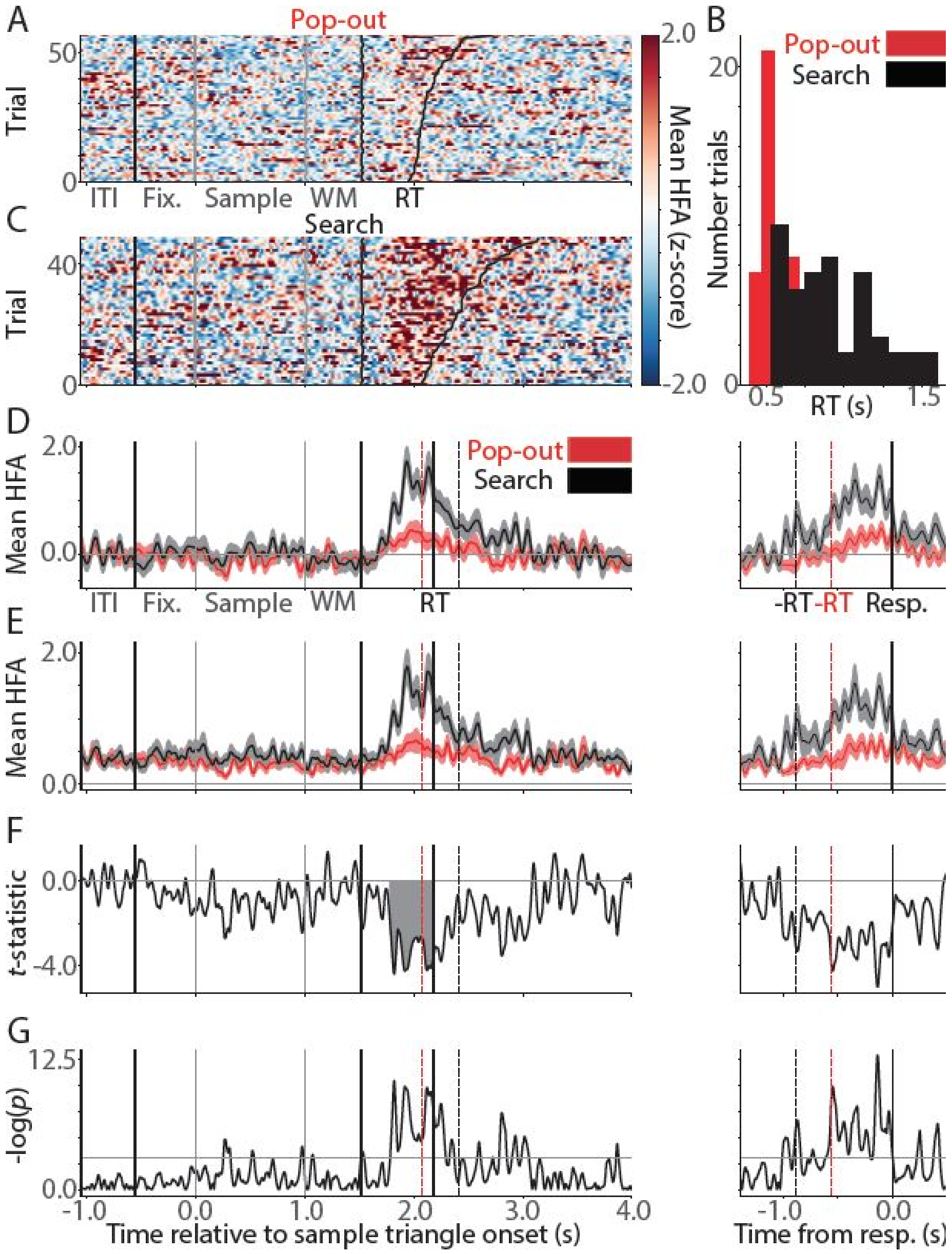


**Supplementary Figure 2.** Example electrode that showed a search-selective effect. A) Stacked single-trial HFA plot for the Pop-out condition, where the trials are ordered by RT so that the fastest trial is on the bottom, and the slowest trial is on the top. Grey labels indicate intervals between vertical lines, similarly to Supplementary Figure 1. ITI, Inter-Trial Interval; Fix., Fixation Interval; Sample, Sample Interval; WM, Working Memory Interval. B) RT distributions for this patient: The RTs can affect the condition-based comparison. C) Stacked single-trial plot for the Search condition. The figure format corresponds to (A). D) Mean HFA activity, z-scored to a pooled baseline, and locked to sample (left) and response (right). The red dashed line on the left panel shows the median RT in Pop-out. The black dashed line shows the median RT in Search. The solid black line between the dashed lines show the median RT across the two conditions. In the right panel, the dashed lines show the median RTs in the two conditions relative to the response (with Pop-out in red and Search in black). E) The same HFA activity is shown as in (D), but here it is zero-clipped, to facilitate comparison between increases only in each condition. F) The two-sample $t$-statistics between the two conditions over time. The shaded area shows a contiguous cluster of significant differences between the two conditions. G) $p$-values corresponding to the $t$-statistics in (F). The horizontal line shows $p = 0.05$.

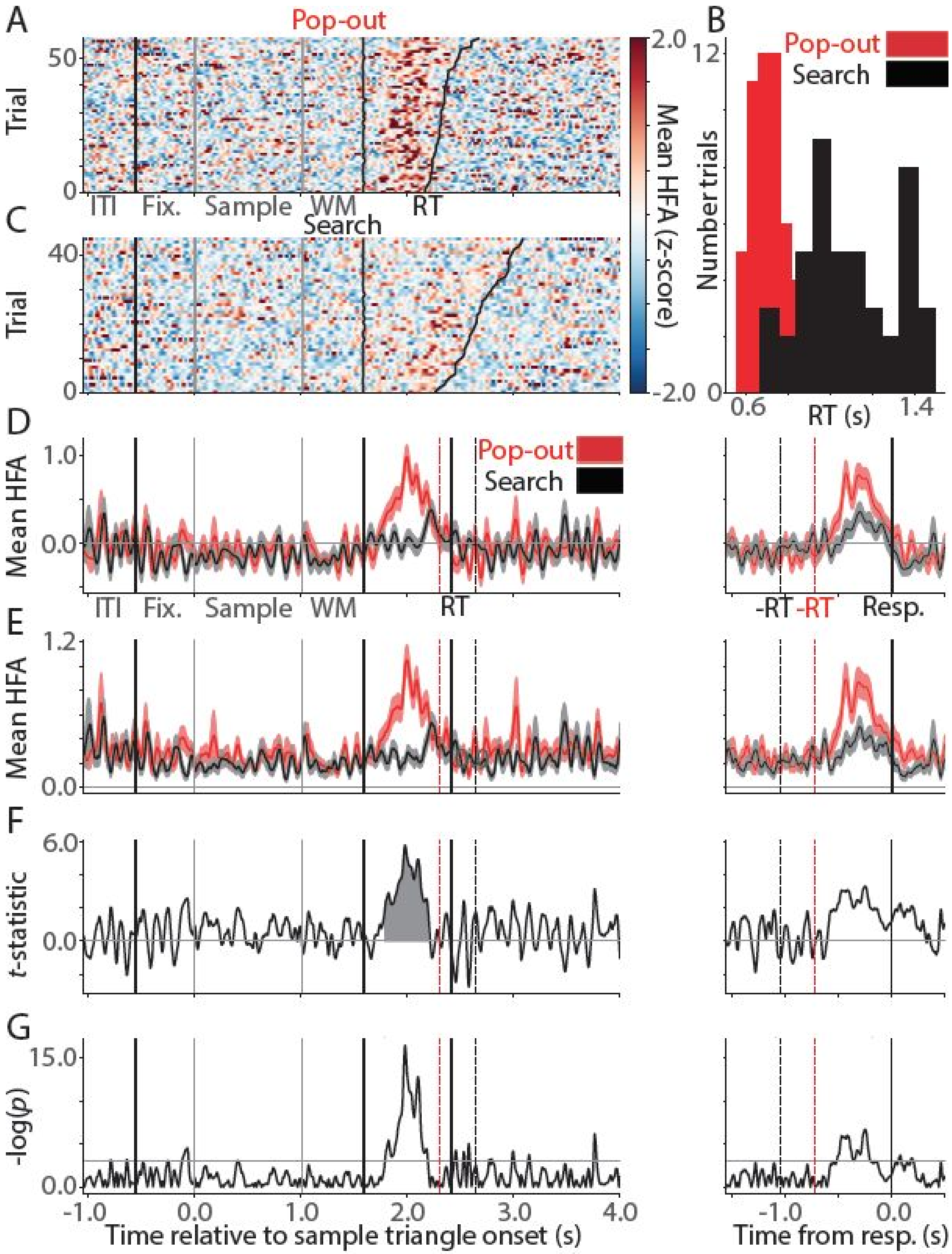


**Supplementary Figure 3.** Example electrode that showed a Pop-out selective effect. The figure layout corresponds to that of Supplementary Figure 2.

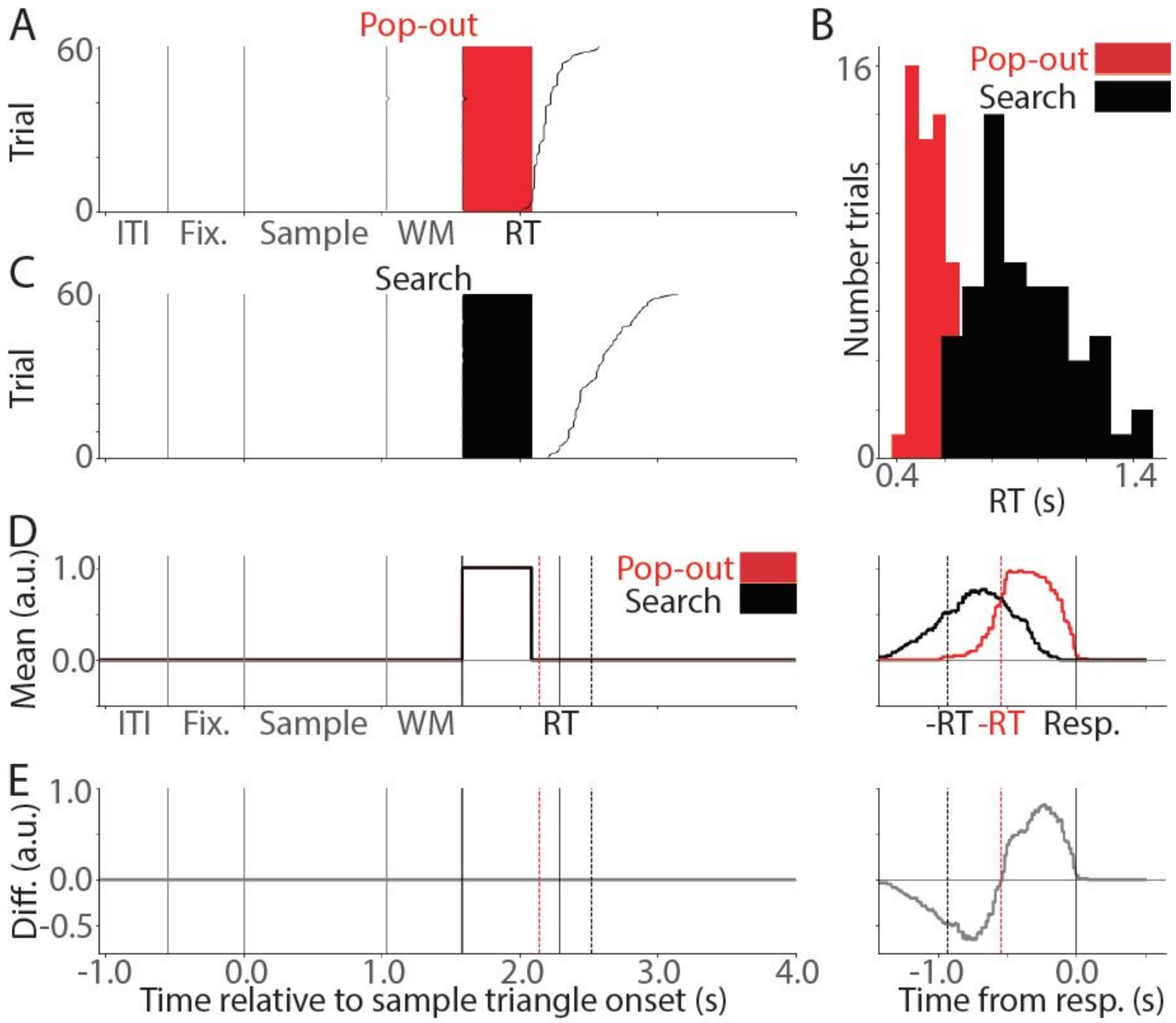


**Supplementary Figure 4.** Simulated electrode showing stimulus-related HFA signal. A) Stacked single-trial plot for the simulated Pop-out condition, comparable to Supplementary Figure 2.A. B) RT distributions for a representative patient, comparable to Supplementary Figure 2.B. These RTs were used to generate the remaining figures in this plot (A, C-E). C) Stacked single-trial plot for the simulated Search condition, comparable to Supplementary Figure 2.C. D) Mean of the single-trial traces, comparable to Supplementary Figure 2.D. The left panel shows stimulus-locked traces, and the right panel shows response-locked traces. a.u., arbitrary units. E) Difference (Diff.) between the mean traces in the two simulated conditions, comparable to the outcome of a two-sample *t*-test as in Supplementary Figure 2.F. The two panels illustrate that, in a stimulus-related electrode in which the two conditions do not differ in their response, the mean signal in the Pop-out and Search conditions will only differ if locked to the response (right panel). This does not affect the outcome of the selection of condition-related effects, which is based on stimulus-locked mean traces.

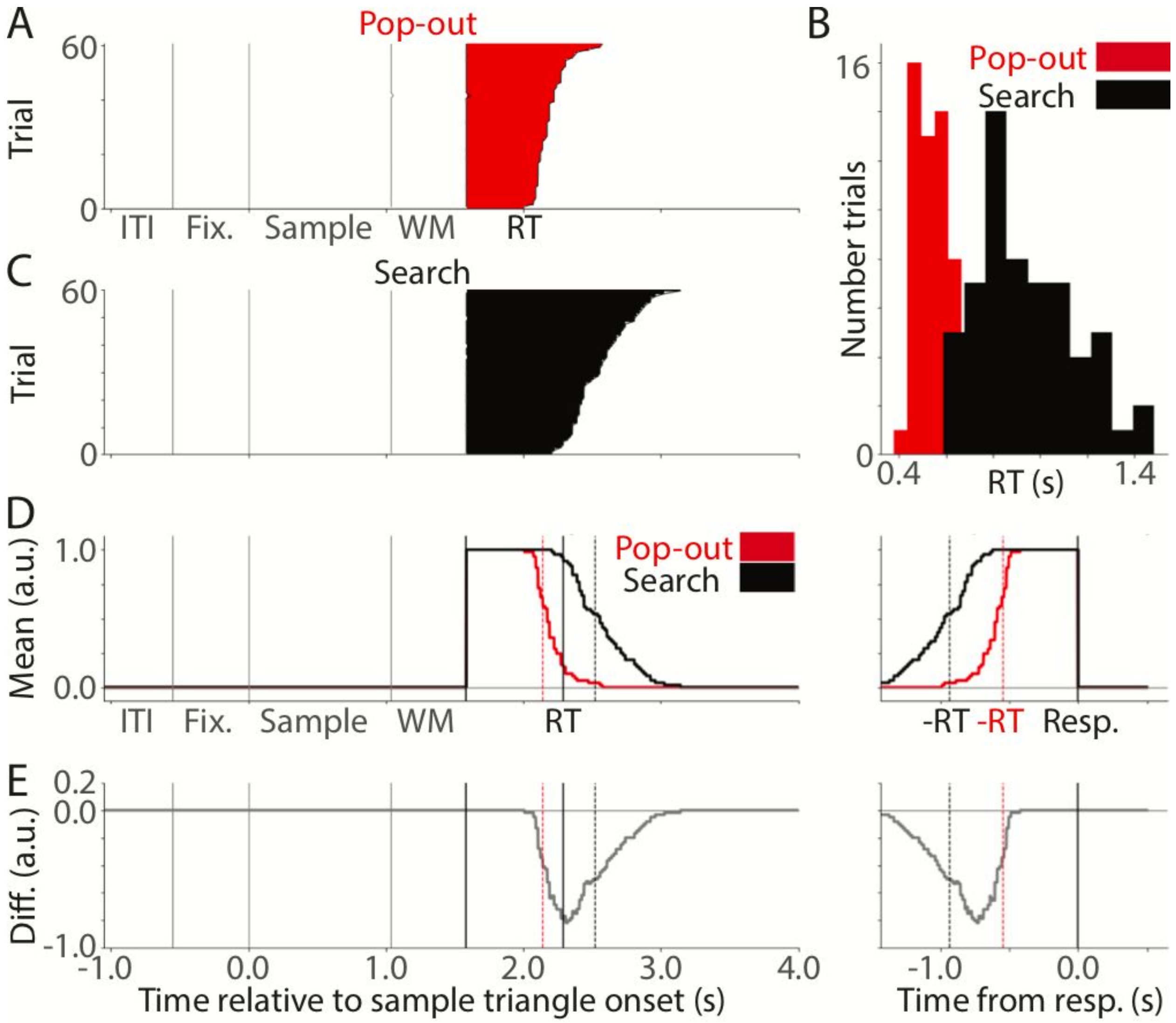


**Supplementary Figure 5.** Simulated electrode showing sustained HFA signal. The figure layout corresponds to that of Supplementary Figure 4. This illustrates that in a sustained-response electrode, in which the simulated response is governed only by the presence or absence of the stimulus screen, the mean signal in the Pop-out and Search conditions will differ such that a spurious Search effect can occur late in the trial interval. Supplementary Figure 7 shows an example from the data, which features this type of artifact.

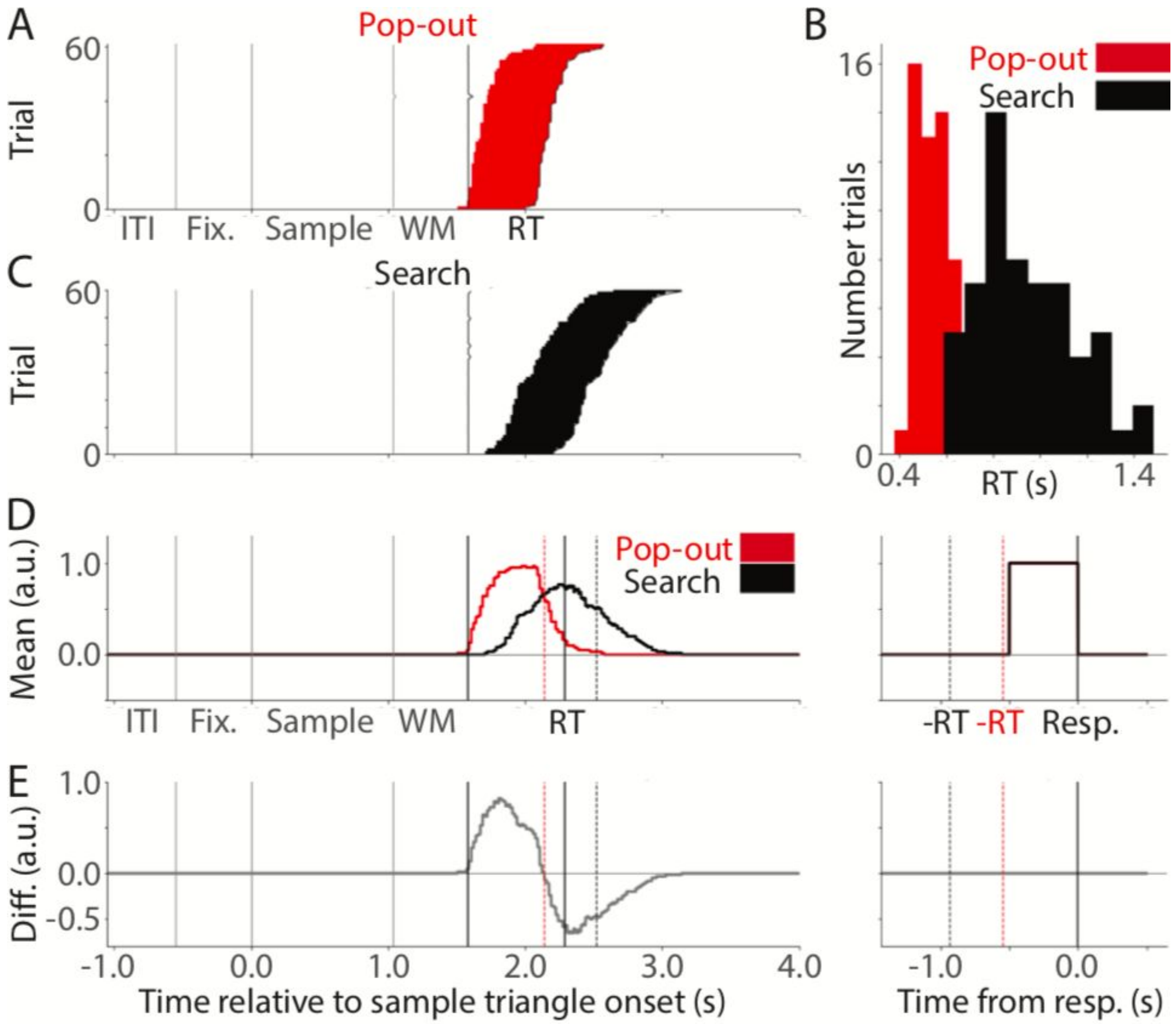


**Supplementary Figure 6.** Simulated electrode showing response-related HFA signal. The figure layout corresponds to that of Supplementary Figure 4. This figure illustrates that, in a response-related electrode, in which the simulated response begins 500 ms before the button press and continues until the time of the button press, the mean signal in the Pop-out and Search conditions will differ such that a spurious Pop-out effect can occur early in the trial interval, while a spurious Search effect can occur late in the trial interval. Supplementary Figures 8 and 9 show examples from the data, in which these types of artifacts occur.

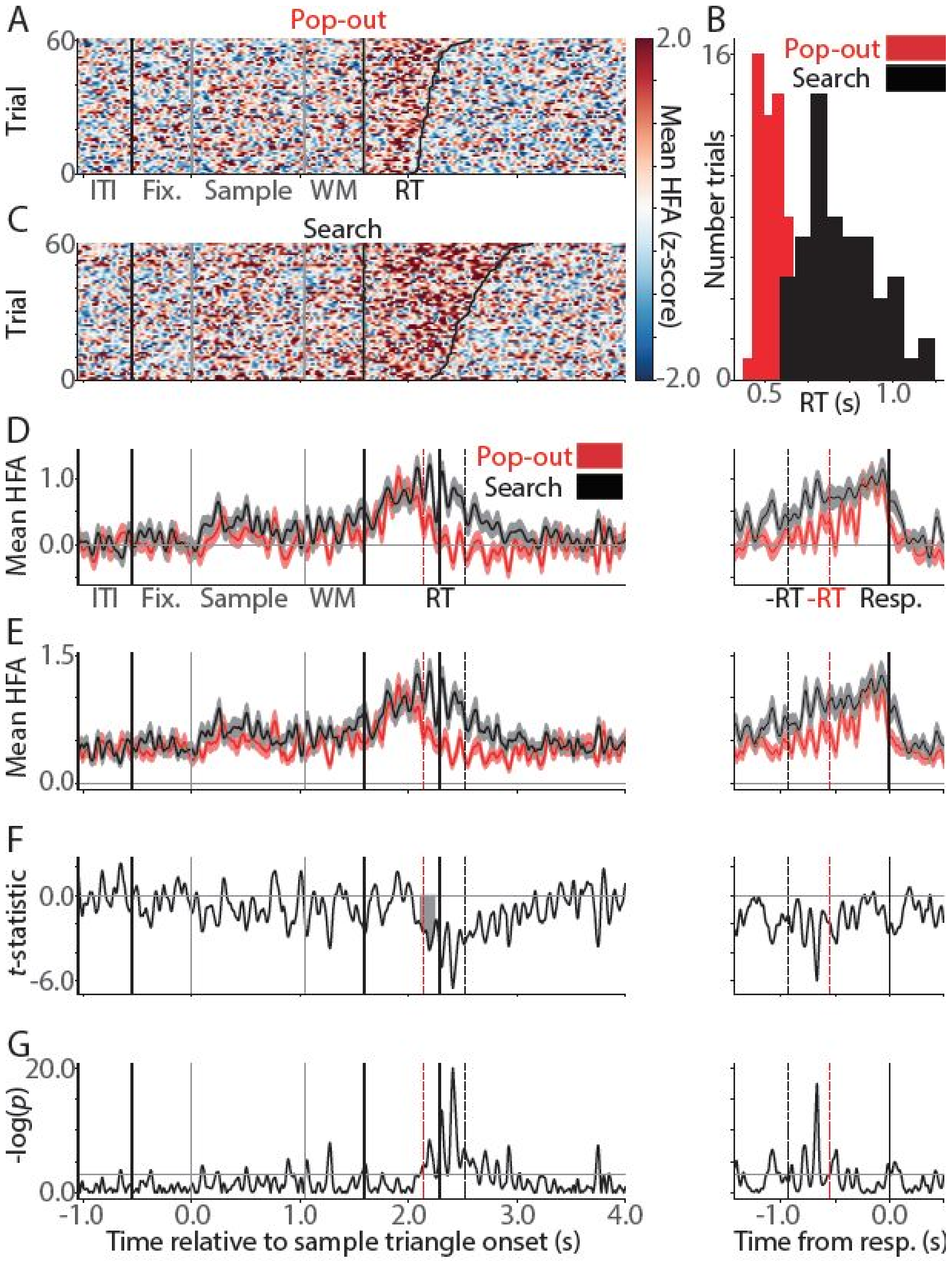


**Supplementary Figure 7.** Electrode with sustained response pattern showing artifactual Search effect. The figure layout corresponds to that of Supplementary Figure 2. This figure shows an example of an electrode, which was automatically identified as showing a Search effect, but where this effect was artifactual due to the sustained response pattern of the electrode and the shape of the RT distribution in this patient. This electrode was labeled as not showing a condition-based effect, based on visual inspection. The observed artifact corresponds to the simulated artifact in Supplementary Figure 5.

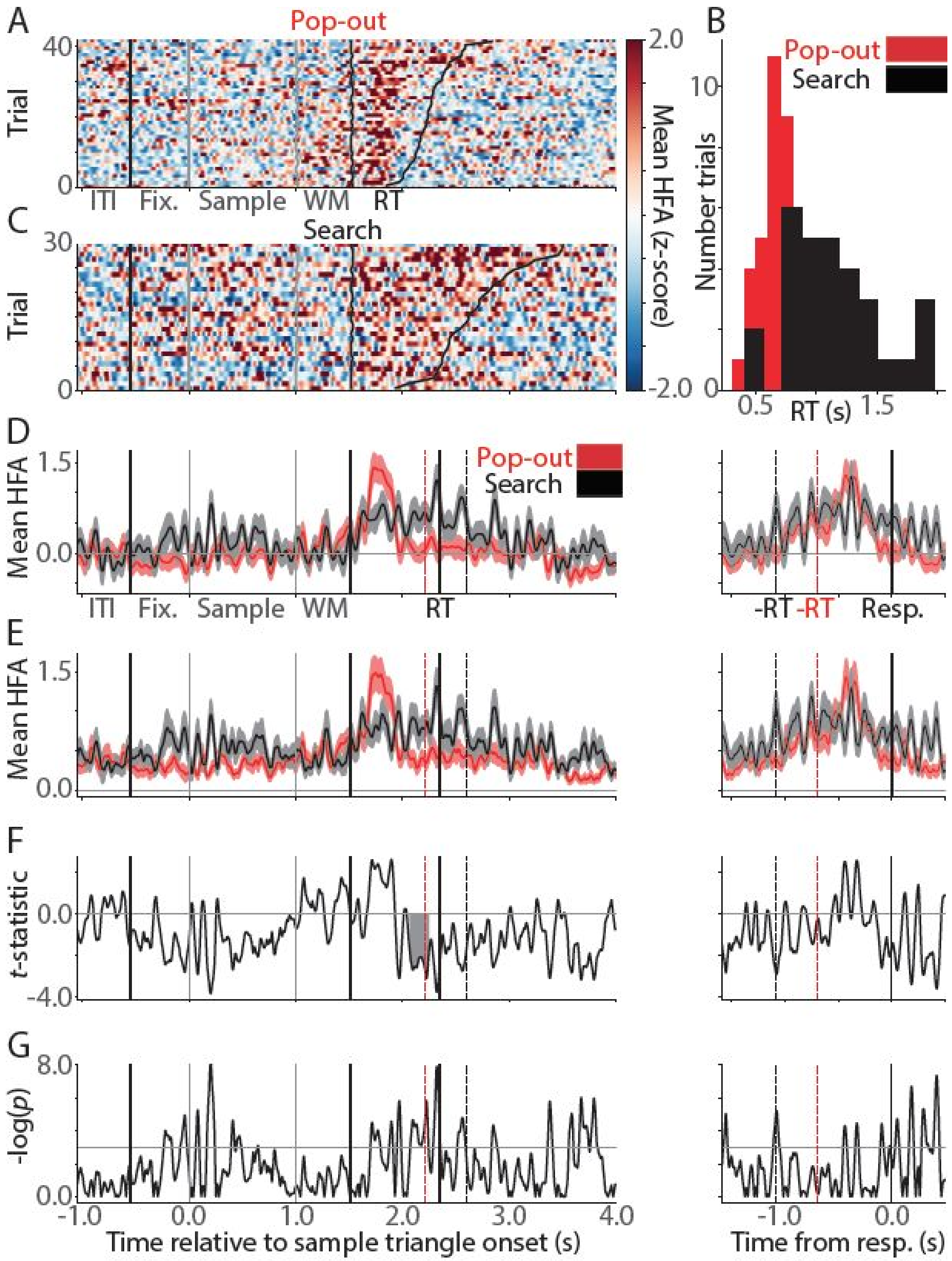


**Supplementary Figure 8.** Electrode with response-related signal pattern showing artifactual Search effect. The figure layout corresponds to that of Supplementary Figure 2. This figure shows an example of an electrode, which was automatically identified as showing a Search effect, but where this effect was artifactual due to the response-related signal pattern of the electrode and the shape of the RT distribution in this patient: The response-locked panels on the right (D-E) show mean traces, which do not differ between conditions. This electrode was labeled as not showing a condition-based effect, based on visual inspection. The observed artifact corresponds to the simulated artifact in Supplementary Figure 6. The shape of the *t*-statistic trace (F) in the present figure can be compared to the condition difference trace in Supplementary Figure 6.

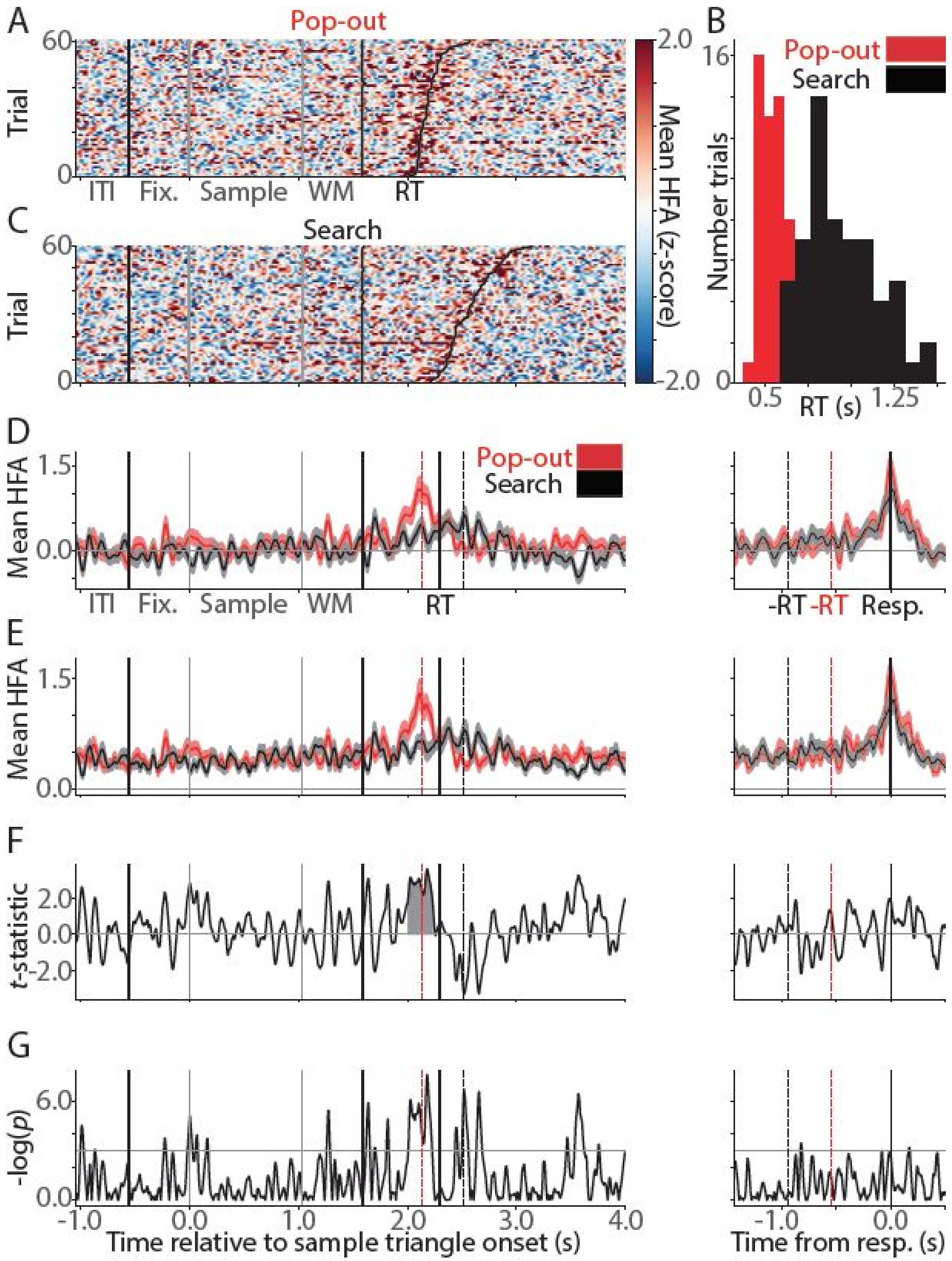


**Supplementary Figure 9.** Electrode with response-related signal pattern showing artifactual Pop-out effect. The figure layout corresponds to that of Supplementary Figure 2. This figure shows an example of an electrode, which was automatically identified as showing a Pop-out effect, but where this effect was artifactual due to the response-related signal pattern of the electrode and the shape of the RT distribution in this patient: The response-locked panels on the right (D-E) show mean traces, which do not differ between conditions. This electrode was labeled as not showing a condition-based effect, based on visual inspection. The observed artifact corresponds to the simulated artifact in Supplementary Figure 6.
